# Inoculation with highly-related mycorrhizal fungal siblings, and their interaction with plant genoptypes, strongly shapes tropical mycorrhizal fungal community structure

**DOI:** 10.1101/2020.07.31.230490

**Authors:** Yuli Marcela Ordoñez, Lucas Villard, Isabel Ceballos, Frédéric G. Masclaux, Alia Rodriguez, Ian R. Sanders

**Author notes:** These authors contributed equally to this work (Yuli Marcela Ordoñez, Lucas Villard). These authors are joint senior authors on this work (Alia Rodriguez, Ian R. Sanders). **Corresponding author:** I. R. Sanders, Department of Ecology and Evolution, University of Lausanne, Biophore building, 1015 Lausanne, Switzerland.

## Abstract

Arbuscular mycorrhizal fungi (AMF) have the potential to increase crop yields and all globally important crops form the mycorrhizal symbiosis. Only a few studies have investigated the impact of introduced AMF on local AMF communities and most studies have only investigated effects of one isolate. We studied the impact on AMF community structure of inoculating roots of the globally important crop cassava with highly genetically-related clonal siblings of two genetically different *Rhizophagus irregularis* isolates. We hypothesized that inoculation with *R. irregularis* siblings differentially influences the structure and the diversity of the pre-existing AMF community colonizing cassava. Alpha and beta taxonomic and phylogenetic AMF diversity were strongly and significantly altered differentially following inoculation with sibling AMF progeny. In most cases, the effects were also cassava-genotype specific. Although biomass production and AMF colonization were also both differentially affected by inoculation with sibling *R. irregularis* progeny these variables were not correlated with changes in the AMF community structure. The results highlight that investigations on the impact of an introduced AMF species, that use only one isolate, are unlikely to be representative of the overall effects of that AMF species and that the genetic identity of the host must be considered. The amount of inoculum added was very small and effects were observed 12 months following inoculation. That such a small amount of almost genetically identical fungal inoculum can strongly differentially influence AMF community structure 12 months following inoculation, indicates that AMF communities in tropical soils are not very resistant to perturbation.

## Introduction

The majority of plants form symbiotic associations with arbuscular mycorrhizal fungi (AMF) in almost all terrestrial ecosystems. These important fungal symbionts, belonging to the subphylum Glomeromycotina, have an enormous potential to increase crop yields on a global scale because they have the potential to improve plant growth and because all globally important crop plants form the mycorrhizal symbiosis [1]. Thus, these fungi could be very important in helping to feed the rapidly growing human population.

Arbuscular mycorrhizal fungi have sometimes proved effective for increasing food production, particularly in the case of cassava; an important food security crop that feeds nearly 1 billion people daily in the tropics [2, 3]. The arbuscular mycorrhizal fungal species *Rhizophagus irregularis* can be produced *in vitro*, thus, free of unwanted microorganisms, and can be highly concentrated into an easy-to-transport product. This fungus is commonly found in agricultural soils around the world, and indistinguishable genotypes of this fungus are distributed across wide geographical, edaphic and climatic areas on different continents, indicating that it is well adapted to many different environmental conditions [4]. To date, only a handful of AMF are able to grow well in an *in vitro* culture system. Application of one *in vitro*-produced isolate of *Rhizophagus irregularis*, resulted in a 20% yield increase in cassava in tropical soils in Colombia [5].

*R. irregularis* has been shown to possess large intra-specific genetic differences [4, 6–9] and genetically different isolates induce differential growth responses in plants [10], including cassava [11]. *R. irregularis* isolates have been shown to exist in two states; as homokaryons, where all nuclei are identical, and as dikaryons, where the fungus contains a population of two nucleus genotypes existing in a common cytoplasm. Production of clonal *in vitro* single spore sibling *R. irregularis* lines (SSSLs), originating from parental dikaryon AMF isolates, were shown to induce enormous variation in rice growth, with some SSSLs greatly increasing rice growth compared to the original parental isolate, even though they are likely to be products of clonal reproduction [12, 13]. Other studies on a dikaryon *R. irregularis* isolate (C3) also showed large differences in quantitative fungal traits among SSSLs [14]. This was thought to be due to the fact that clonal offspring spores originating from a dikaryon AMF parent can inherit unequal proportions of two nucleus genotypes [15]. Because of the inter-isolate and within-isolate genetic variation and its concurrent effects on plant growth, plus the fact that it can be grown *in vitro*, *R. irregularis* has been considered a potentially good candidate for an AMF improvement program [16]. Furthermore, more is known about the genome of this AMF species than any other AMF species. Such a use of improved *R. irregularis* lines would represent a fundamental change to the way that AMF inoculants are used [11].

The study by Angelard et al. 2010 [12] on variation in rice growth with AMF was conducted in sterile soil. However, in farming situations, AMF have to be applied to soil that naturally already contains an existing diverse AMF community and a diverse microbiome [1], meaning that while variation among SSSLs may induce variation in plant growth in sterile conditions, it may not translate into such impressive effects in the field. Therefore, two field trials were conducted in Colombia in order to investigate the potential of variation among *R. irregularis* SSSLs, on cassava production [11]. Two different varieties of cassava were each inoculated with two *Rhizophagus irregularis* lines, representing two genetically different parental isolates, known as C2 and C3. Isolate C2 is a homokaron and C3 is a dikaryon [17]. The plants were also inoculated with 3 SSSLs of parental isolate C2 and 9 SSSLs of the parental isolate C3 [11]. The *R. irregularis* SSSLs originating from both parents induced large significant differences in cassava root production, with up to three-fold differences among plants inoculated with *R. irregularis* SSSLs originating from the same parent [11]. Isolate C2 and its progeny induced differential growth in both cassava cultivars but in a different way in each cultivar and isolate C3 and its progeny only induced differential growth in one cultivar (CM4574). The differential effects of the SSSLs lines was also dependent on the cassava cultivar. The effect of the *R. irregularis* lines on cassava production was reproducible in two trials in separate years [11].

Commercial AMF inoculants are often applied to soils from which those fungi were not isolated. Because the application of AMF inoculants to crops represents the addition of an AMF isolate to a pre-existing AMF community, it is important to understand the effects of this practice on the existing AMF community [18]. However, there is very limited empirical data from field inoculation trials on the effect of an introduced AMF on the diversity and structure of the local AMF community, especially in trials where AMF inoculation treatments actually led to strongly significant effects on crop growth. In two studies in temperate soils there was no effect on the existing AMF community of adding one *R. irregularis* isolate [19, 20]. In experimental microcosms with synthetic communities and field trials with a local AMF community, inoculation with *R. irregularis* supressed most of the AMF taxa [21–23] and decreased AMF diversity [24]. All the studies used only one fungal isolate of an AMF species as inoculum.

The study by Ceballos et al. (2019) [11] is the only field trial to date showing significant and very large variation in the yields of a crop following inoculation with single spore progeny derived from homokaryon and dikaron AMF parents. Thus, there is currently no knowledge about whether variation among clonally produced single spore progeny could also alter local AMF communities.

If inoculation with SSSLs lines results in large variation in root growth, there is a strong rationale for expecting that a change in the local AMF community would occur. We propose that if more resources were allocated by the plants below-ground, following inoculation with a given SSSL, then this could also have significantly affected resource availability to the AMF community in the roots, as well as potentially affecting root exudation, and how many nutrients were removed from the soil. Thus, it is likely that the AMF in the different SSSL treatments experienced differential resource availability. It is well known that changes in resource availability greatly influence community diversity and structure [25, 26]. Thus, different levels of resources available to AMF, or the competitive ability between the introduced and pre-existing AMF taxa, could cause changes in local AMF community diversity and structure. We, therefore, expected that inoculation with SSSLs that induce differential growth responses in cassava will induce changes in the AMF community.

In first trial conducted by Ceballos et al 2019 we, therefore, investigated the effects of inoculation with *R. irregularis* SSSLs on the structure and diversity of the AMF community inside the roots of two varieties of field grown cassava in Colombia [11]. We did not attempt to track the inoculated fungal isolates in the field because genetically very similar *R. irregularis*-like fungi already existed in the soil at the field site, thus, making it extremely difficult to distinguish those that were introduced from those that were already present. Given the observed effects of the SSSLs on variation in cassava root growth, we hypothesized that inoculation of cassava with *R. irregularis* SSSLs influences the structure and/or the diversity of the pre-existing AMF community. Secondly, we hypothesized that the changes observed in the AMF community would be more pronounced in the cassava variety that showed a greater variation in growth response to the different *R. irregularis* SSSLs and where variation among SSSLs induced different levels of AMF colonization in the roots.

## Materials and methods

### Field site

The field experiment was conducted at the Utopía campus of La Salle University, Yopal, Department of Casanare, Colombia (72° 17’ 48 W, 5° 19’ 31” N). The climate is tropical with average temperatures of 18°C (night) to 28°C (day), with an average air humidity of 75% and total annual precipitation of 2335 mm with 172 rain days. Physical and chemical soil properties are shown in Table S1.

### Plant and fungal material

We used cassava cultivars MCOL2737 and CM4574. These cultivars were selected because they are suitable for production in that region of Colombia and because MCOL2737 was used in previous experiments [5, 27].

We used two isolates of *R. irregularis* (C2 and C3) that we also refer to as parental isolates. The two fungi originate from an agricultural field in Switzerland and were subsequently put into *in vitro* culture in 2000 [6]. They have subsequently been maintained at the University of Lausanne. Each isolate was originally initiated with one spore. The SSSLs of C2 and C3 were raised *in vitro* in 2008 and then maintained in the same conditions by sub-culturing roots, hyphae and spores every 3-4 months. The SSSLs of parental C2 are referred to as C2.1, C2.2. and C2.3. The SSSLs of parental C3 are referred to as C3.1, C3.2, C3.3, C3.4, C3.5, C3.6, C3.7, C3.8 and C3.9. The SSSLs of C2 and C3 appear genetically identical to the parental fungus qualitatively, meaning that they possess the same alleles as the parent. However, some of the SSSLs derived from the C3 parental isolate differ from each other in allele frequency at bi-allelic loci [15].

### Experimental design

The field experiment was established in a randomized factorial design with nine replicate blocks, 15 different inoculation treatments (14 different *R. irregularis* lines and one mock-inoculated treatment) and two cassava cultivars as factors. There was one replicate of each treatment combination per block and 9 replicates per treatment combination. To avoid cross contamination between treatments in the field, experimental units (inoculated or mock-inoculated plants) were positioned 3 metres apart, with two non-treated plants at one metre distances in between. Cassava was planted in rows, with a planting density of 10000 plants ha^−1^, which is the standard planting density recommended in the region. Plants were given fertilizer at the rate of 233 Kg ha^−1^ urea, 125 Kg ha^−1^ di-ammonium phosphate (DAP), 100 Kg ha^−1^ potassium chloride (KCl) and 41 Kg ha^−1^ of Kieserite (a fertilizer comprising 3% soluble potassium, 24% magnesium and 19% sulphur) and 22 Kg ha^−1^ of Vicor (a granular fertilizer comprising 15% nitrogen, 5% calcium, 3% magnesium, 2% sulphur, 0.02% boron, 0.02% copper, 0.02% manganese, 0.005% molybdenum, 2.5% zinc). The level of P is 50% of that recommended for the region. This was used because previous experiments showed that there was no significant increase in cassava root yields in the same soil using 100% of the recommended P application compared to 50% P application [5]. Half of the fertilizer was applied 45 days after planting and the rest was applied 90 days after planting.

Cassava stem cuttings were inoculated with 500 propagules of *R. irregularis*, at 15 and again at 45 days after planting. Each plant was inoculated by one *R. irregularis* line. The mock-inoculated treatment received the same volume of water.

### Sampling, harvesting and measurements

Three replicate plants of each treatment combination were tagged at the start of the experiment. Three months after inoculation, fine roots of tagged plants were sampled using a soil corer, washed and stored at −80°C for later molecular analysis.

Twelve months after inoculation, each inoculated or mock-inoculated plant was removed by hand. Bulbous roots were weighed and then dried and weighed again. Root weight data are publicly available in a separate manuscript [11]. Fine roots were treated in the same way as the root samples taken 3 months after inoculation, except that a sub-sample of roots was stained to measure colonization by AMF (as described in Ceballos et al. 2019). Because it is often difficult to recover fine cassava roots, we were not able to obtain root samples of all replicates. The numbers of replicates sampled is shown in Table S2.

### DNA extraction and sequencing

One hundred mg of each root sample was used to extract DNA with the PowerSoil DNA isolation kit (MoBio Laboratories Inc., Carlsbad, CA, USA). The small ribosomal subunit (SSU) region was amplified from each DNA sample using the primer pair AML1 and AML2 [28]. The PCR was carried out in a final volume of 20 ml, using 0.1 mM dNTPs, 10 pmol of each primer, 5U of Taq polymerase and the supplied reaction buffer (Thermo Fisher Scientific), under the following conditions: initial denaturation at 94°C for 3 min, 30 cycles at 94°C for 30 s, 50°C for 1 min, 72 °C for 1 min, and a final extension period at 72°C for 10 min. The first PCR product was then diluted 1/100 with distilled deionized H2O. The dilutions were immediately used in a second PCR reaction performed using the NS31 and AM1 [29, 30] with the same PCR regime, except that the annealing temperature was 58°C. Primer NS31* had a 5 nt extension used as a barcode to distinguish sequences from the different samples after sequencing.

Following the second PCR step, amplicons of around 550 bp were separated in an agarose gel and excised. The DNA was extracted with a gel purification kit (Wizard^®^ SV Gel and PCR Clean-Up System, Promega). Amplicons were pooled and the concentration was standardized. Then libraries were constructed using the Illumina TruSeq^®^ DNA Sample Preparation high-throughput (HT) kit. Library quality was assessed by capillary electrophoresis through a Fragment Analyzer™ (Advanced Analytical Technologies, Inc.). In total, 13 libraries were made with a mean number of pooled samples per library of 34 (range 17-45). Sequencing of the libraries was carried out on a 300 PE MiSeq platform. The 13 libraries were distributed in 9 lanes for sequencing. Some libraries were run individually in one lane and some other libraries were run with another library in the same lane. Pooling more than two libraries in a lane did not affect OTU number.

### Bioinformatic analysis

Illumina adapter sequences were removed from the reads, as well as nucleotides having an average Phred score below 15 within a 9bp window. Reads containing undetermined nucleotides (i.e. N) were discarded. All these steps were performed with TagCleaner and PrinSeq. Good quality reads were then checked for correct barcodes, demultiplexed and merged using PEAR software by specifying a minimum merged read length of 200bp. At this stage, 27520013 reads with a mean length of 511bp were retained. Primer and barcode sequences were then removed and reads with a length below 500bp were discarded.

Remaining reads were trimmed to a 500bp length and sorted according to their nucleotide sequence (hereafter referred to as rRNA allele) and their relative abundance in the dataset. This left a total of 15883170 reads that were analyzed as the whole dataset comprising all treatments and both sampling times. Alleles that were present in only one root sample were considered uninformative for AMF community comparisons and were, therefore, removed from the dataset. This represented 1’405’160 alleles following read trimming and singleton removal. Of these, 4’936’382 sequences were submitted to the DBC454 software. This corresponds to the collection of alleles weighted by their relative occurrence in the dataset (i.e. an allele found in 100 root samples occurred 100 times in the dataset). The alleles were submitted to a BLAST search against INSDC (http://www.insdc.org) and MaarjAM (https://maarjam.botany.ut.ee) databases with default parameters in order to determine their putative taxonomic origin. Each allele was classified as either non-AMF, potential AMF or valid AMF, depending on the identity of the best match in both databases and the percentage of similarity to it. We used a 95% similarity threshold to the INSDC database and a 97% similarity threshold to MaarjAM database, to distinguish valid AMF from potential AMF. We also assigned a putative AMF family and virtual taxa (VT) to each allele considered as either potential AMF or valid AMF by using the identity of the best match given in the MaarjAM database.

### OTU model selection

OTUs were assigned by a hierarchical clustering model using DBC454 v1.4.5 [31]. Because this procedure differs from the commonly used VT classification system, the selected models were compared to a VT classification using the Adjusted Rand Index [32]. Because model selection is essential for optimizing the numbers of OTUs that are reliably detected, we provide full details of the pipeline used for model selection as a Supplementary note 1.

### Phylogeny

Phylogenetic relationships between valid AMF OTUs from the selected DBC454 model were inferred by building a phylogenetic tree, using consensus sequences based on nucleotide frequency for each OTU, as well as SSU rRNA type sequences from known AMF species determined by [33]. Sequences were aligned using MAFFT 7.058. The phylogenetic tree was inferred using MrBayes 3.2.2 [34] with a GTR+G nucleotide substitution model and *Schizosaccharomyces pombe* SSU rRNA sequence as the tree root. Two independent runs of MCMC sampling for 2 million generations were performed with a sampling of chain state at each 1000 generations. A maximum clade credibility tree was compiled after checking for run convergence and discarding 30 percent of trees as a burn-in fraction.

### Patterns of OTU community diversity (alpha diversity)

The Shannon-Weiner Index was used to determine OTU taxonomic alpha diversity in each sample. Phylogenetic alpha diversity was assessed by calculating the net-relatedness index (NRI). Replicates were then pooled by AMF treatment and cassava cultivar treatments. We used the “mock-inoculated” treatment to determine the null community mean and standard deviation of diversity indices for each cassava variety independently. Significant differences between null community diversity and *R. irregularis* treatments for each cassava variety were assessed using ANOVA and a Tukey honest significant difference (HSD) post-hoc test. We also assessed the relationship between OTU diversity in each sample and cassava root dry-weight as well as AMF root colonization level using the Pearson’s product moment correlation coefficient.

### Patterns of OTU community turnover (beta-diversity)

In order to investigate whether OTU taxonomic turnover and phylogenetic turnover was affected by *R. irregularis* line and cassava cultivar treatments, we built pairwise dissimilarity distance matrices between each AMF treatment of a given cassava variety by using the Bray-Curtis distance and mean nearest taxon distance (MNTD), respectively. Significant effects of *R. irregularis* lines on both metrics of community turnover were assessed using PERMANOVA analysis. We also ran hierarchical clustering methods using *pvclust* in R 3.4.2 (http://www.R-project.org) [35]. The “average” method was implemented in *pvclust* and we assessed cluster robustness with a bootstrap resampling strategy.

### OTU spatial autocorrelation

Even though the experimental design should limit the bias of OTU distribution due to dispersal limitation, we investigated whether spatial distance between samples in the field site could explain the observed OTU distribution patterns. Therefore, we performed Mantel tests between community diversity indices, as well as inferred distance matrices of community turnover, and spatial distances matrices to test whether alpha or beta diversity correlated with spatial distances among plants.

### Effect of sampling time on alpha diversity and community turnover

Differences in alpha taxonomic and phylogenetic diversity among the same plants that were sampled at 3, and again at 12 months post-inoculation were tested using paired *t*-tests. Differences in community taxonomic and phylogenetic turnover between the two sampling times were assessed by performing PERMANOVA analyses on the Bray-Curtis and MNTD, with sampling time as a factor. Hierarchical clustering was also performed on the data at the two time periods.

## Results

The pipeline, and results of the clustering model, the parameters determined to give optimal OTU selection and the characteristics of the dataset used for the subsequent analyses are all presented in Supplementary note 1. The placement of the OTUs inside the phylogenetic tree built with SSU reference sequences was coherent with their taxonomic classification using best match annotation of BLAST results (Figure S1). The vast majority of OTUs fell into the Glomeraceae clade (44 OTUs; 96.95% of reads), and mostly within the *Rhizophagus* genus. The second most abundant group was in the Acaulosporaceae family (6 OTUs, 2.9% of reads), followed by Claroideoglomeraceae (2 OTUs, 0.084% of reads) and Gigasporaceae/Scutellosporaceae (2 OTUs, 0.051% of reads).

### Colonization by AMF

Inoculation with SSSL progeny of isolate C3 significantly affected the levels of colonization by AMF inside the cassava roots but the way the SSSLs affected AMF colonization was not the same in the two cassava cultivars (Figure S2). Colonization by AMF did not differ significantly among plants inoculated with C2 and its SSSL progeny in either cultivar (Figure S2). Colonization by AMF differed significantly among cassava roots that were inoculated with the parental isolate C3 and its progeny SSSLs. However, the differences among treatments were not the same in the two cassava cultivars. Colonization was lowest in plants inoculated with parental isolate C3 and highest in plants inoculated with the SSSL C3.6 in cultivar CM4574. In cultivar MCOL2737, AMF colonization was highest in plants inoculated with SSSL C3.3 and lowest in plants inoculated with SSSL C3.7.

### Alpha diversity - Shannon-Weiner Index

Twelve months following inoculation, AMF taxonomic alpha diversity (as measured by the Shannon-Weiner index) was significantly affected by the AMF inoculation treatments and by cultivar but with no significant interaction between these two factors (Table S3). A further ANOVA performed separately on the data of each cultivar showed that alpha diversity was only affected by the *R. irregularis* treatments in cultivar MCOL2737 (Table S4). In this cultivar, alpha diversity was significantly the highest in roots of cassava inoculated with *R. irregularis* SSSL C3.6. Alpha diversity in these plants differed significantly from alpha diversity in the mock-inoculated, C3.1 and C3.9 treatments (Figure 1). Lines C3.6 (highest value) and C3.9 (lowest value) were SSSLs derived clonally from the same parental isolate.

**Figure 1:**
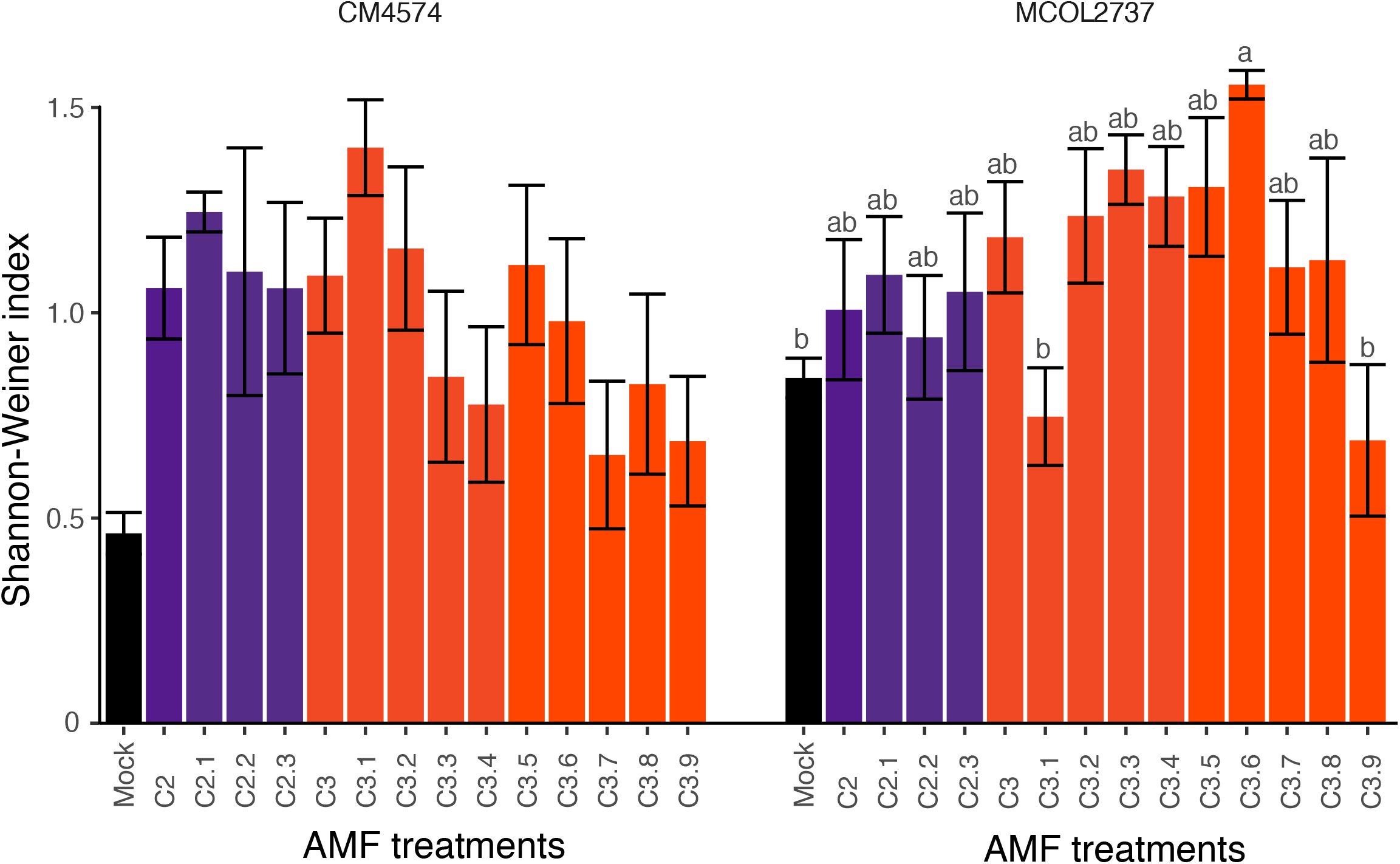
Shannon-Weiner index of OTU diversity inside the roots of cassava varieties CM4574 and MCOL2737 twelve months following inoculation with the AMF line treatments. Bars in purple represent treatments with parental isolate C2 and its corresponding SSL offspring. Bars in orange represent treatments with parental isolate C3 and its corresponding SSL offspring. Mock corresponds to the mock-inoculated treatment. Any bars with the same letter above means that they are not significantly different according to a Tukey HSD post-hoc test (*P* < 0.001).

There was no significant correlation between the Shannon-Weiner index, root colonization and dry cassava weight. The Shannon-Weiner index was related to the number of reads and the number of OTUs in each root sample (Table S5). No correlation between differences in diversity and spatial distance was observed (Table S5).

### Alpha diversity - phylogenetic diversity (NRI)

*R. irregularis* line treatments strongly affected the NRI of AMF communities in cassava roots, while there was no significant effect of cassava cultivar (Table S3). A marginally significant interaction between the two factors was observed (Table S3). Similarly to the Shannon-Weiner index, a further ANOVA performed separately on the datasets for each cassava cultivar showed that significant differences in NRI were only observed in the roots of cultivar MCOL2737 (Table S4). The mean pairwise distance (MPD) is calculated on a presence-absence matrix with a larger value representing closer phylogenetic distance and smaller values indicating greater phylogenetic distance among the members of the AMF community. The phylogenetic distance among the AMF community in the mock-inoculated treatment was the highest suggesting a close phylogenetic similarity among the AMF taxa in the pre-existing AMF community (Figure 2). In contrast, AMF communities in cultivar MCOL2737 inoculated with parental isolate C2 and its SSSLs and parental isolate C3 showed a significantly smaller MPD value indicating inoculation with those treatments resulted in more phylogenetically diverse AMF communities in the roots (Figure 2).

**Figure 2:**
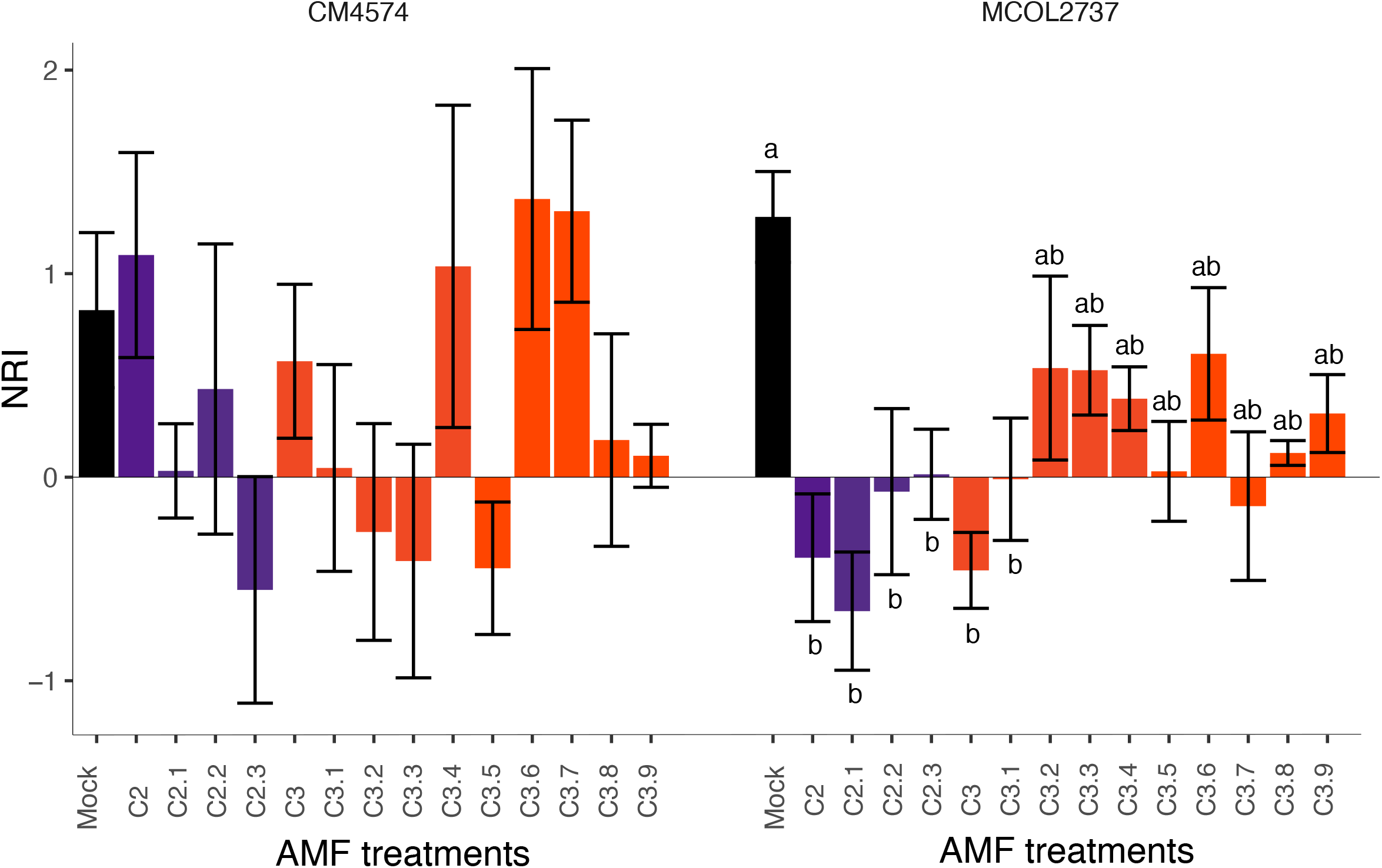
Net relatedness index (NRI) of OTUs inside roots of cassava varieties CM4574 and MCOL2737 twelve months following inoculation with the AMF line treatments. Bars in purple represent treatments with parental isolate C2 and its corresponding SSL offspring. Bars in orange represent treatments with parental isolate C3 and its corresponding SSL offspring. Mock corresponds to the mock-inoculated treatment. Any bars with the same letter above means that they are not significantly different according to a Tukey HSD post-hoc test (*P* < 0.001).

We also found a negative relationship between the Shannon Weiner index and NRI, which meant that as AMF communities became more taxonomically diverse they also became more phylogenetically diverse (Figure S3).

### Beta diversity- taxonomic and phylogenetic turnover

Bray-Curtis dissimilarity indices of the AMF communities differed significantly among *R. irregularis* line treatments in both cultivars although the way this measurement was affected by the different AMF treatments differed significantly between the two cassava cultivars. (Tables S6 and S7). We performed a hierarchical analysis based on Bray-Curtis dissimilarity matrices that was conducted on data from each cassava cultivar separately. A significant cluster represents a group of two or more cassava plants whose roots contained taxonomically similar AMF communities. The clustering showed that the taxonomic composition of the AMF communities in both cultivars was clustered significantly at 12 months after inoculation with different fungal lines but that different patterns of clustering were observed in the two cultivars (Figures 3a and 3b) meaning that the cassava cultivar also determined the taxonomic composition of the AMF communities.

**Figure 3:**
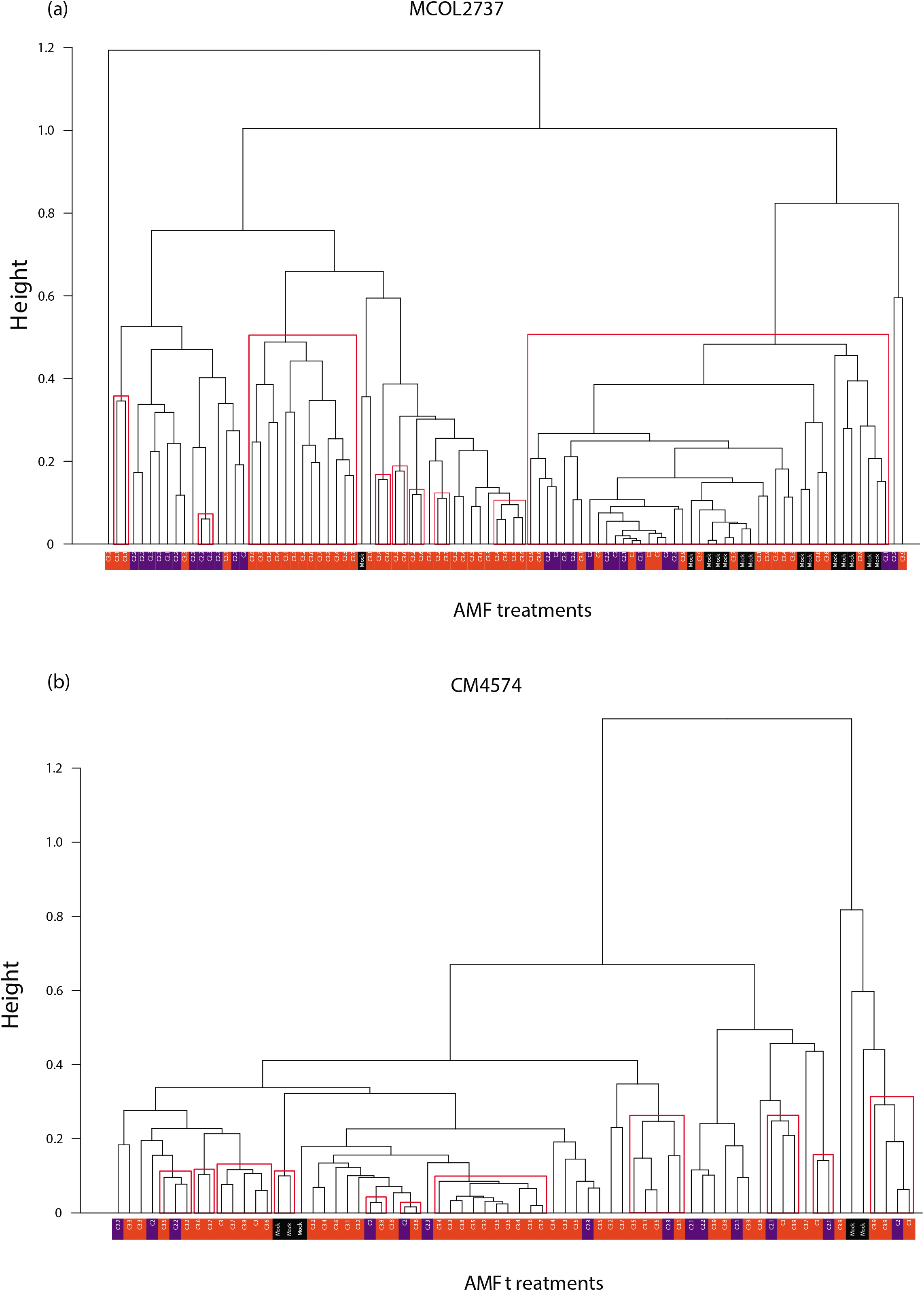
Hierarchical clustering based on Bray-Curtis Dissimilarity in **(a)** cassava variety MCOL2737 **(b)** cassava variety CM4574. Red squares indicate >95% probability of the existence of a cluster using the approximated-unbiased value. Bars in purple represent treatments with parental isolate C2 and its corresponding SSL offspring. Bars in orange represent treatments with parental isolate C3 and its corresponding SSL offspring. Mock corresponds to the mock-inoculated treatment.

Phylogenetic turnover, or beta phylogenetic diversity, was assessed by calculating the mean-nearest taxon distance (MNTD). Significant differences in the MNTD of AMF communities among AMF line treatments occurred as well as between the two cultivars (Table S6). Because there was also a significant AMF treatment x cassava cultivar interaction we also analysed the effects of AMF inoculation treatments on each cassava cultivar separately, showing that there was a significant effect in cultivar MCOL2737 but not in cultivar CM4574 (Table S6 and Table S7). Hierarchical clustering based on MNTD also showed more discrete clusters of phylogenetically similar AMF communities occurred in cassava variety MCOL2737 (Fig. S4a) compared to cultivar CM4574 (Fig. S4b).

### Differences in alpha and beta diversity at 3 and 12 months following inoculation

In order to investigate possible changes in AMF community diversity in roots between 3 and 12 months, we analysed paired data from individual tagged plants, sampled at 3 and again at 12 months following inoculation. A paired *t*-test was then used to compare values at 3 and 12 months from the same plants. Results showed that AMF community alpha diversity, measured as the Shannon-Weiner index, was higher at 12 months than at 3 months (Figure 4a, Table S8), even though there was no increase in read number or OTU number during this time period in those plants (Table S8; Figures S5 and S6). Mean phylogenetic diversity indices were also influenced by the number of months following inoculation with a strong decrease in phylogenetic relatedness in the AMF communities between 3 and 12 months following inoculation (Table S8; Figure 4b).

**Figure 4:**
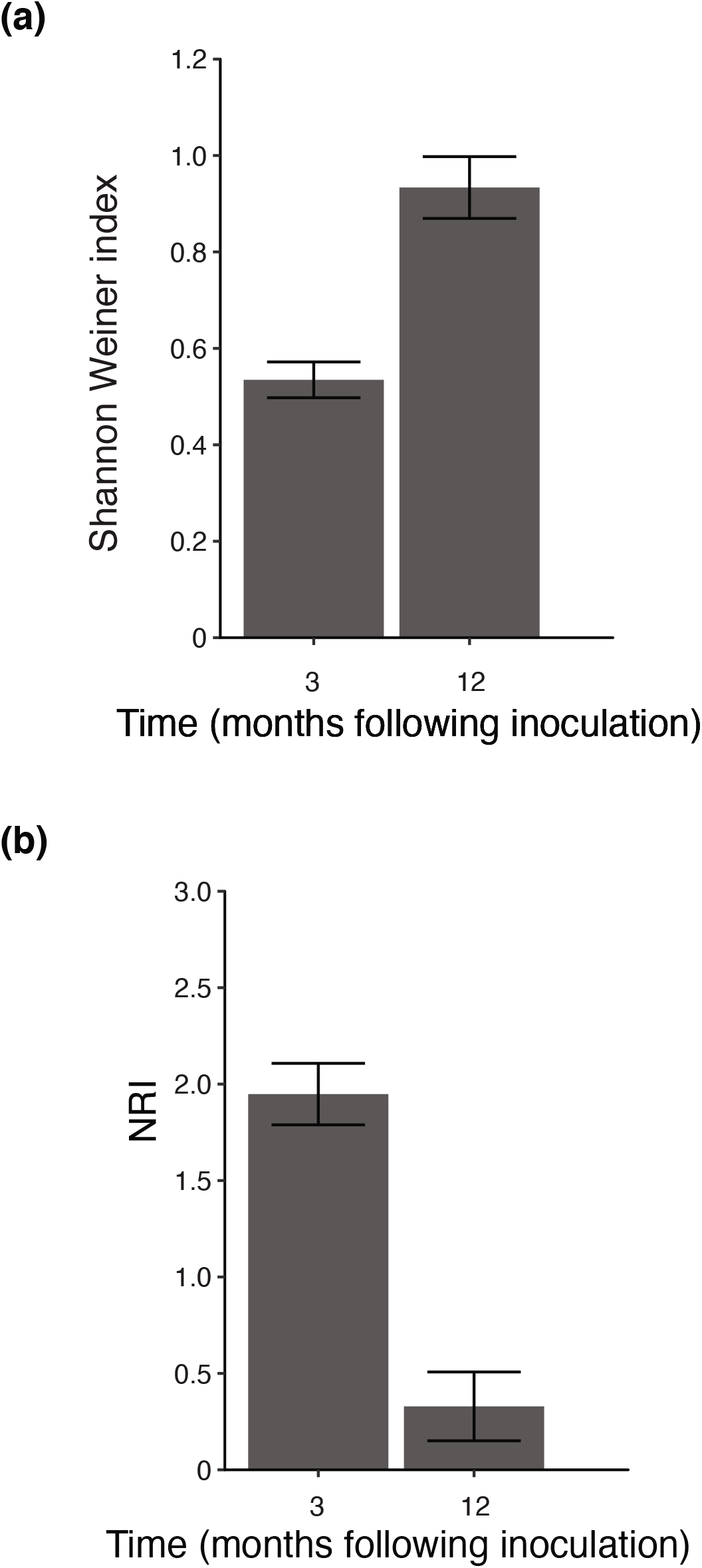
**(a)** Bar-Plot showing Shannon Weiner index measured in the same individual plants sampled at 3 and 12 months following inoculation. See Table S8 for results of paired t-test. **(b)** Bar-Plot showing NRI measured in the same individual plants sampled at 3 and 12 months following inoculation. See Table S8 for results of paired t-test.

AMF beta diversity at both the taxonomic and phylogenetic level, as measured by the Bray-Curtis Distance and MNTD, showed that the similarity in the AMF community composition (as shown by hierarchical clusters of the data) changed between 3 and 12 months following inoculation in MCOL2737 (Figures 5 and 6; Table S9). Taxonomic composition of the AMF communities was more similar among plants at 3 months, as shown by larger clusters of similarity compared to 12 months following inoculation (Figure 5), indicating a change in AMF taxonomic community composition over time. Phylogenetic beta diversity in MCOL2737 roots also showed that clusters of plants showing communities with similar composition of species with a given phylogenetic relatedness occurred at both 3 and 12 months following inoculation. However, the clusters were not comprised of the same treatments at 3 and 12 months showing that differences phylogenetic composition of the community changed over time.

**Figure 5:**
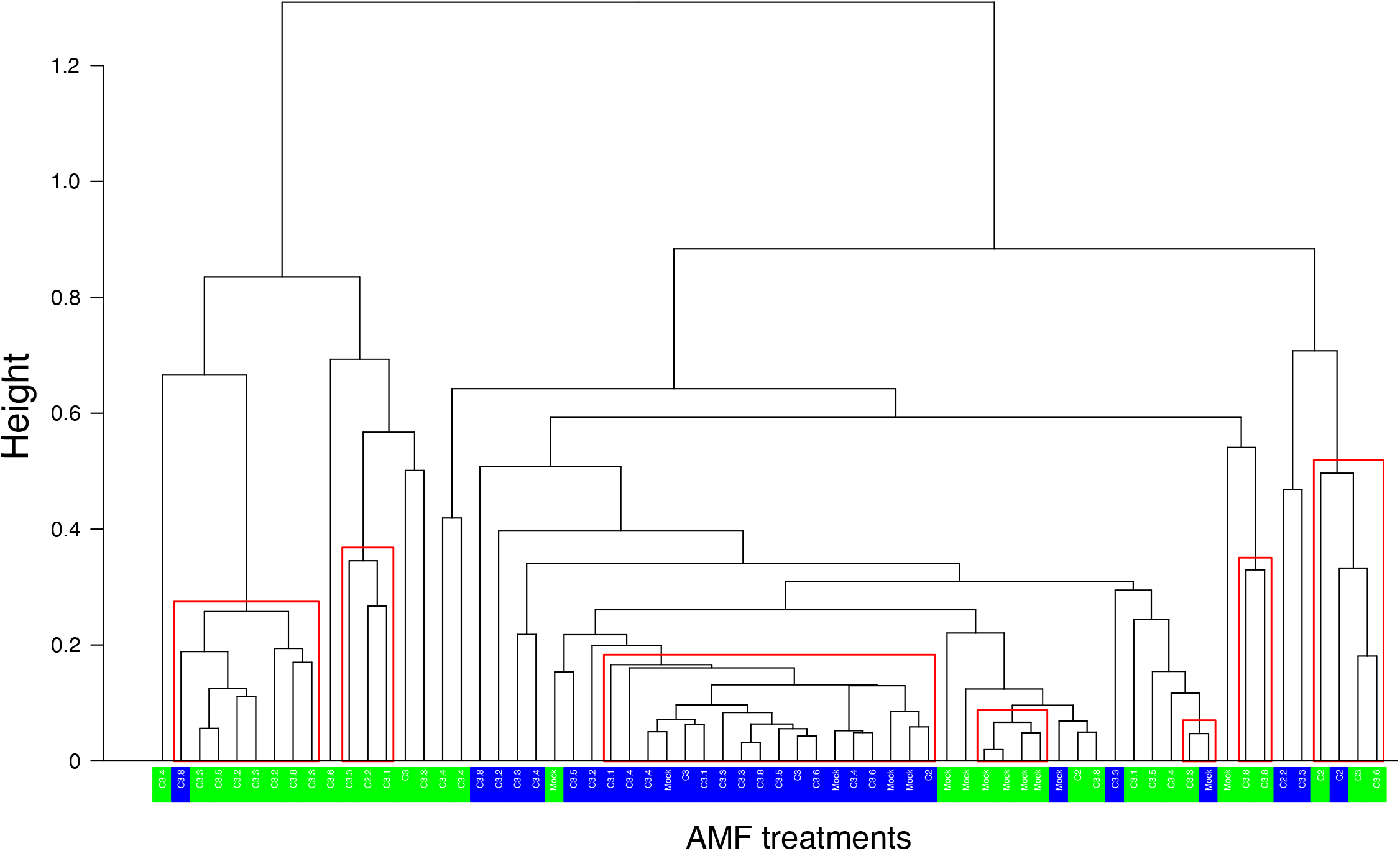
Hierarchical clustering based on Bray-Curtis Dissimilarity index for paired cassava plants of cultivar MCOL2737 sampled at 3 months (in blue) and 12 months (in green) following inoculation. Red squares indicate a >95% probability of the existence of a cluster using the approximate-unbiased value. Mock corresponds to the mock-inoculated treatment.

**Figure 6:**
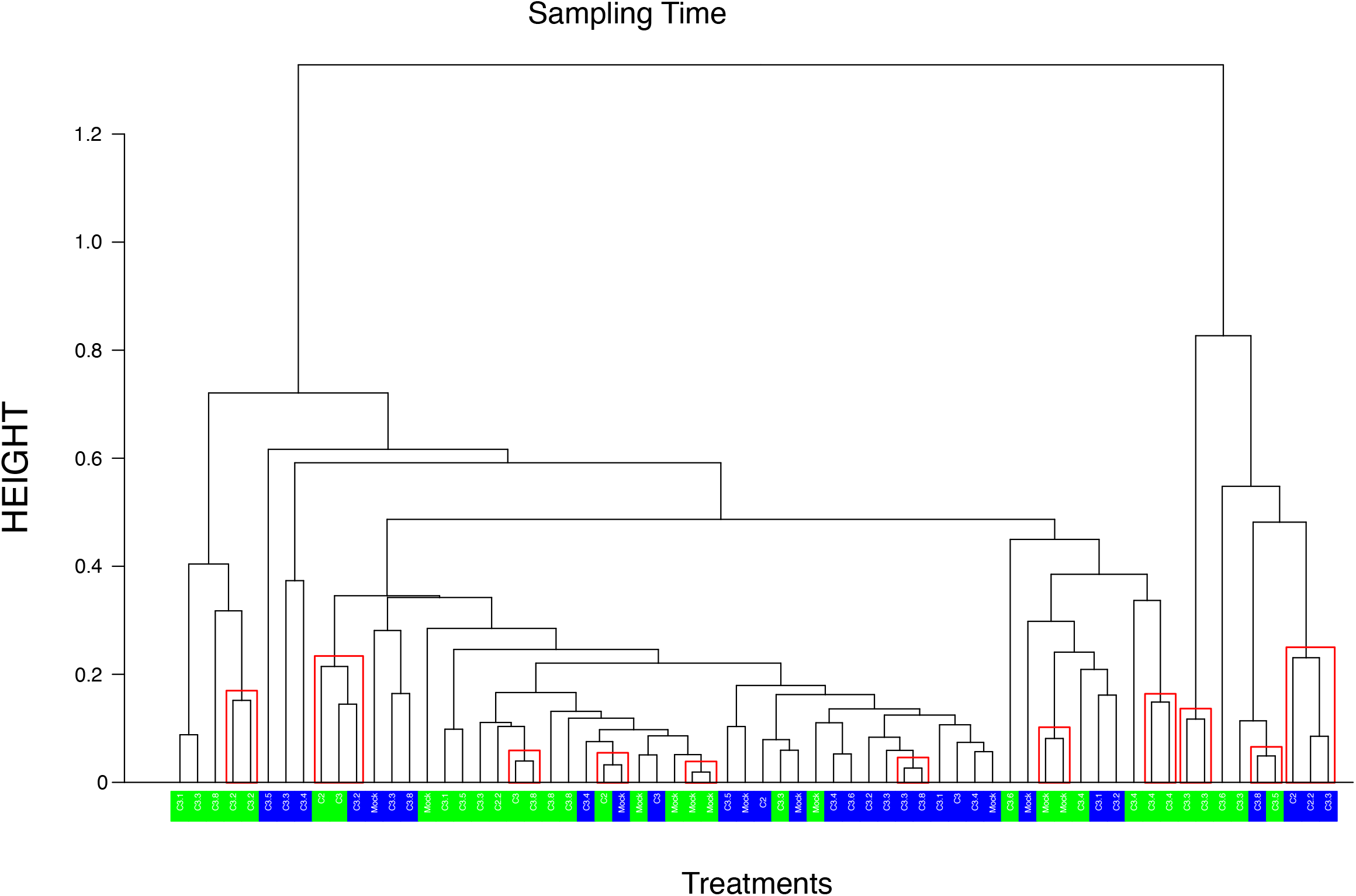
Hierarchical clustering based on Mean Nearest Taxon Distance (MNTD) for paired cassava plants of cultivar MCOL2737 sampled at 3 months (in blue) and 12 months (in green) following inoculation. Red squares indicate a >95% probability of the existence of a cluster using the approximate-unbiased value. Mock corresponds to the mock-inoculated treatment.

## Discussion

While there are many descriptive studies looking at AMF diversity and community structure in different vegetation types, soil types and environmental gradients, there are surprisingly few studies looking at AMF inoculation effects on the local AMF community in the field and in an experimental framework. Our study is unique in that we experimentally studied the effect of genetically different parental isolates of the same AMF species, *R. irregularis*, and clonal SSSLs produced by each of the parental fungi on existing AMF communities. We tested the effects of these inoculation treatments on the AMF communities in two cassava cultivars that differed strongly in their growth responses to inoculation and in which inoculation also affected overall AMF colonization. The SSSLs are clonal offspring of the parents and yet surprisingly strong differential effects on the local AMF community were observed.

### AMF inoculation and its effect on local AMF community diversity

We found that adding clonal progeny of the same species of AMF, *R. irregularis*, had surprisingly large and significant effects on both alpha and beta diversity of AMF communities at both the taxonomic and phylogenetic levels but that this was also highly dependent on the variety of the plant. The results are in strong contrast to most of the published studies of effects of *R. irregularis* on the local AMF community. The effects appeared largely independent of the strongly differential effects of inoculation on cassava root growth (observed by Ceballos et al. 2019) [11] or overall levels of AMF colonization by the AMF community. Inoculating cassava with a parental *R. irregularis* isolate and with its clonal single spore offspring did not significantly reduce the diversity of the AMF community or supress colonization by other AMF species. In fact, in some cases the AMF taxonomic alpha diversity increased in the roots of cassava in response to inoculation compared to the mock inoculated treatment and inoculation with some other siblings of the same parental fungus. Furthermore, inoculation with some of the fungi resulted in AMF communities that were phylogenetically more diverse compared to the mock inoculated treatment. The effects of inoculation with the different fungal treatments not only occurred on levels of alpha diversity but the AMF community composition (beta diversity) was also affected by inoculation with parental and SSSL progeny. All effects on alpha and beta diversity were strongly cassava cultivar dependent.

### Surprising ecological effects of clonal SSSLs on AMF communities

As stated in the hypotheses, we expected to see effects of inoculation on the AMF community because we had observed that inoculation with SSSL progeny of an AMF parent strongly influenced cassava roots growth and because the clonal SSSLs also altered overall AMF colonization. However, we consider a number of effects of inoculation with these fungi to be very surprising that we discuss below.

Firstly, a diverse AMF and microbial community normally exists in soils. We added an extremely small amount of fungal material (approximately 1000 spores) to cassava stakes at planting and in a soil with an existing diverse microbiome. The majority of the effects were measured 12 months following inoculation. This means that despite adding only a very small amount of a fungal material, this had profound and long-lasting influence on levels of taxonomic and phylogenetic diversity as well as the composition of the AMF community.

Secondly, the fungal inocula were clonal progeny of two different parental genotypes of the same fungal species, *R. irregularis*. Isolate C3 is a dikaryon. Progeny of this isolate, produced clonally *in vitro*, share identical alleles. However, allele frequencies at bi-allelic sites differ in their proportion among progeny, representing differences in the frequency of the two nucleus genotypes [15]. This means that each sibling contains the same genetic information but at some loci, the proportions of two alleles can differ. We find it surprising that such small quantitative genetic differences among siblings of the same parent can induce such large differences in their effects on the local AMF community. These siblings also strongly differentially affected how much the AMF community colonized the roots. Parental isolate C2 is a homokaryon. This means that all offspring are genetically identical. We did not see differential effects among C2 and its siblings on alpha diversity. While the C2 family did not have significant effects on taxonomic alpha diversity, all these isolates influenced the AMF community in the same way by increasing the phylogenetic diversity among members of the AMF community compared to the control.

Taken together, these results mean that adding a very small amount of very highly related fungal material, with what would be considered as representing miniscule genetic differences, to a pre-existing diverse AMF community can have long lasting effects on the AMF community at least one year later. Most significantly, this means that AMF communities in tropical agricultural soils appear to be extremely sensitive to perturbations, even to tiny differences in the inoculum.

### Are inoculation effects on the AMF community linked to plant growth or colonization?

We saw significant effects of inoculation with *R. irregularis* SSSLs on the diversity and structure of AMF community. In this sense, the results partially support the hypotheses. However, we hypothesized that an alteration in AMF communities would occur where we observed effects of inoculation on plant growth and AMF colonization. Changes in cassava root growth and overall levels of AMF colonization of the roots would be a good indication of an alteration in availability of resources for AMF. In a diverse AMF community, we would expect that different AMF taxa would respond differently to an alteration in resource availability, thus driving a change in the AMF community. While, we observed changes in the AMF community, these were not obviously related to overall changes in root growth or overall levels of AMF colonization.

### Plant genotype specific effects on AMF communities

One notable result of this study is that in most cases, the differential effects on the AMF community, and on AMF colonization, induced by SSSLs of *R. irregularis* were strongly cassava cultivar dependent. Differential effects of the *R. irregularis* SSSLs on cassava growth in Colombia, Kenya and Tanzania were also strongly cultivar dependent [11]. What these results mean is that inoculation of crops with AMF can be altered significantly using inoculum with very small genetic differences but that the effect of inoculation on the local AMF community will likely be different according to which cassava variety the farmer chooses. Given the strong cultivar effects, a detailed investigation into the relationship between cassava genetic variability and its influence on AMF communities is highly warranted. At present, in crop production, the choice of cultivar is primarily considered before the choice of whether to inoculate with AMF. In the case of cassava cultivation, cultivar choice also determines the effects of inoculation and the subsequent consequences on the local AMF community.

### Time effects

The comparison of the AMF community between 3 and 12 months following inoculation clearly indicated that taxonomic alpha diversity greatly increased over this period while the community became composed of more distantly related taxa. Thus, the AMF community that colonizes cassava roots is dynamic. We specifically focused our main harvesting effort at 12 months because previous studies showed that AMF inoculation effects on cassava are usually seen in the last month of the cassava growth period [2, 5] when the roots are rapidly accumulating starch, but we cannot rule out that earlier effects on the AMF community that were not measured in this study may have been correlated with inoculation effects on cassava root weight at the final harvest.

### Conclusions

The central question of this study addresses an important ecological issue regarding how sensitive local AMF communities are to perturbation by very closely related AMF within the same species. This study indicates that even addition of very small amounts of highly related AMF can perturb the local AMF community in tropical soils and in a way that is also highly plant genotype specific. While other studies have tested one AMF isolate on the local AMF community, with contrasting results, we show that even very small genetic changes in the inoculum can give completely contrasting effects ranging from no effect on the local AMF community to a large change in AMF alpha and beta diversity in the local community. Our study provides a framework for specifically testing a series of new and pertinent ecological questions, namely: 1. Why does an AMF community respond differently to additional of highly related fungi? 2. Are alterations to the AMF community that are caused by inoculation with an AMF detrimental or beneficial? 3. In the case where inoculation causes changes in plant growth but does not cause changes in the AMF community, are inoculation effects direct or mediated through other components of the soil microbiota? While researchers have put a lot of effort into describing AMF communities using DNA sequencing approaches, at present, there is a great lack of experimental studies looking at which factors can alter AMF diversity and community structure. We strongly encourage community ecologists interested in AMF community diversity to focus on more experimental studies that can help to unravel these important ecological questions about the use of AMF inoculants.

## Supporting information

Supplementary note 1: Pipeline for OTU model selection

## Acknowledgments

We thank the La Salle University for allowing us to conduct the field trial at the Utopia campus, the Utopia students for helping establish and maintain the experiment and Ricardo Peña Venegas and Cristhian Fernandez for logistical support. We thank the International Center for Tropical Agriculture (CIAT) for providing cultivars CM4574 and University La Salle for providing cultivar MCOL2737. All bioinformatics computations were performed at the Vital-IT (www.vital-it.ch) Center for High Performance Computing of the Swiss Institute of Bioinformatics and all Illumina MiSeq^®^ sequencing was performed at the Genomic Technologies Facility (GTF) of the University of Lausanne. This work was supported by two Swiss National Science Foundation grants to IRS (grant numbers: IZ70Z0-131311/1 and 31003A_162549) and a COLCIENCIAS scholarship to YMO and IC. Sequence reads were deposited in the NCBI SRA database (BioProject Accession Number: SRP133761).

## Conflict of interest

The authors declare no conflict of interest

## Supplementary Material

Supplementary information is available with this manuscript, including 6 supplementary figures and Supplementary note 1.

## Notes

### Competing Interest Statement

The authors have declared no competing interest.

