## Supplementary figures S1-S6 for "Inoculation with highly-related mycorrhizal fungal siblings, and their interaction with plant genoptypes, strongly shapes tropical mycorrhizal fungal community structure"

**Figure S1**: Maximum clade credibility phylogenetic tree of consensus sequences aligned with a selection of reference sequences from Krueger et al.(2012). OTUs in blue represent those found in this study.

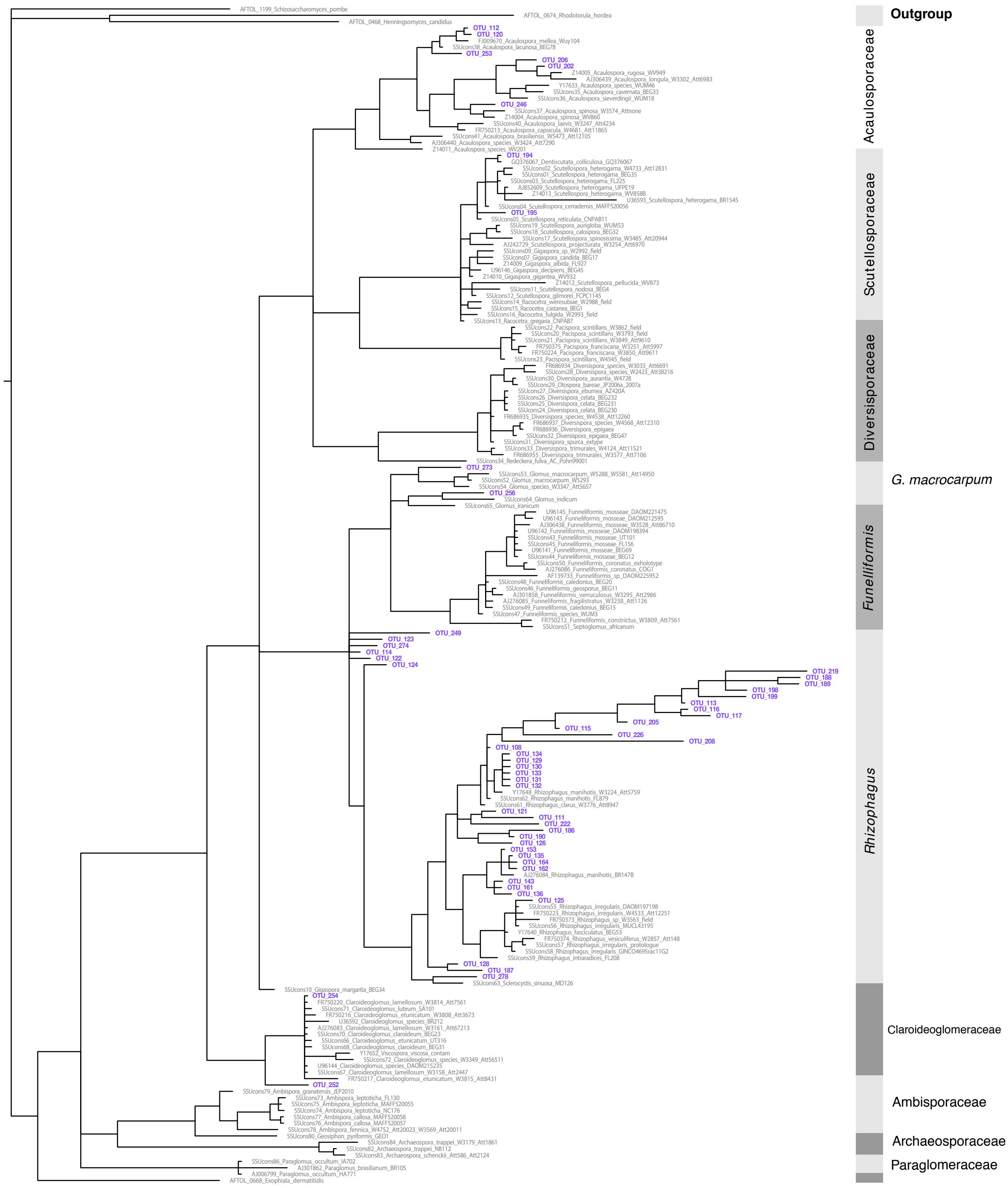

**Figure S2**: Colonisation by AMF inside the roots of cassava varieties CM4574 and MCOL2737 twelve months following inoculation with the AMF treatments. Purple bars represent inoculation with parental isolate C2 or its SSSL offspring. Orange bars represent inoculation with parental isolate C3 or its SSSL offspring. Black bars correspond to the mock-inoculated treatment. There was a significant main AMF treatment effect on the whole dataset and a strongly significant AMF treatment x cassava variety interaction. The interaction terms were significant with or without including the mock treatment at $P \leq 0.0001$. Because of the strongly significant interaction, the dataset was split and re-analysed by AMF family (parental isolate and SSSL siblings) and cassava variety. This analysis revealed that there were no significant differences in AMF colonisation among plants inoculated with C2 and its progeny in either cassava variety. There were strongly significant differences in AMF colonization among plants inoculated with parental isolate C3 and its progeny in both cassava varieties (at $P \leq 0.0001$ and $P = 0.0272$, in varieties CM4574 and MCOL2737, respectively). Bars with the same letter above means that they are not significantly different according to a Tukey HSD post-hoc test conducted separately for each cassava variety ($P < 0.001$).

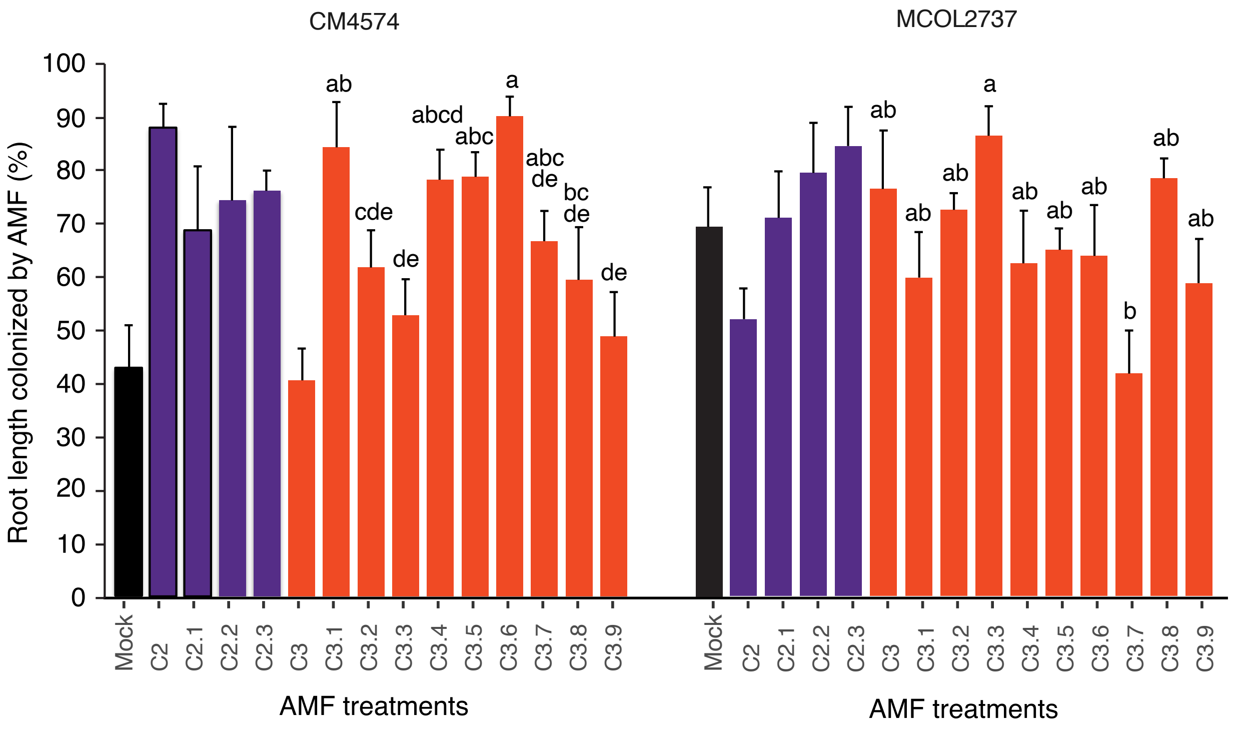

**Figure S3**: Relationship between Shannon-Weiner index and Nearest Relative Index (NRI) of AMF communities from cassava roots sampled 12 months following inoculation. The relationship is significant (y=-0.7x, p-value<0.01)

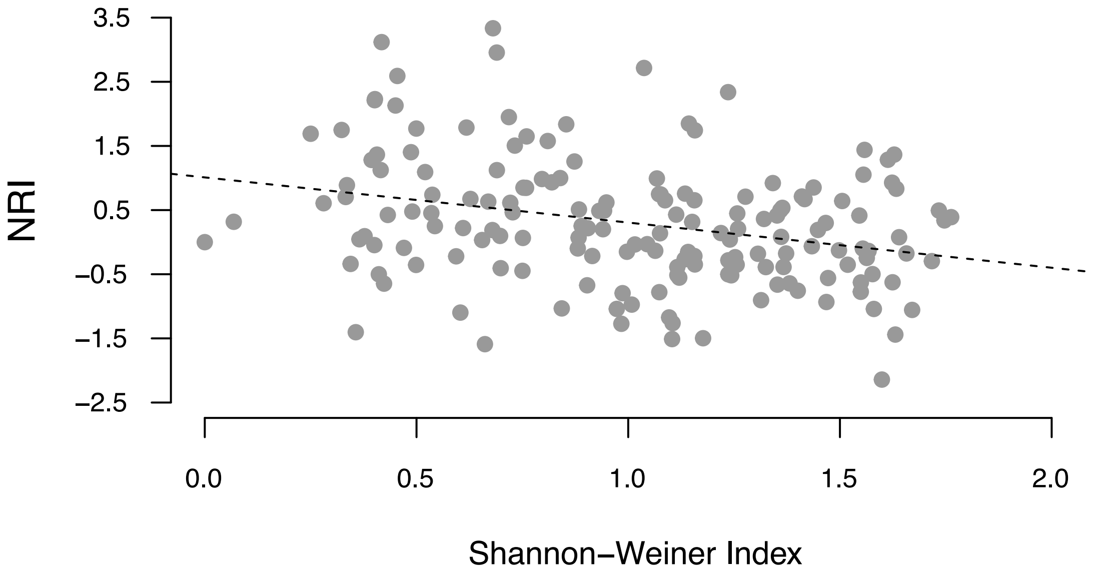

**Figure S4:** Hierarchical clustering based on Mean Nearest Taxon Distance (MNTD) in **(a)** cultivar MCOL2737 **(b)** cultivar CM4574. Red squares indicate >95% probability of cluster existence using the approximatively-unbiaised value. AMF treatments with the same colour represent a parental AMF isolate and its corresponding SSL offspring. Mock corresponds to the mock-inoculated treatment.

**a.**

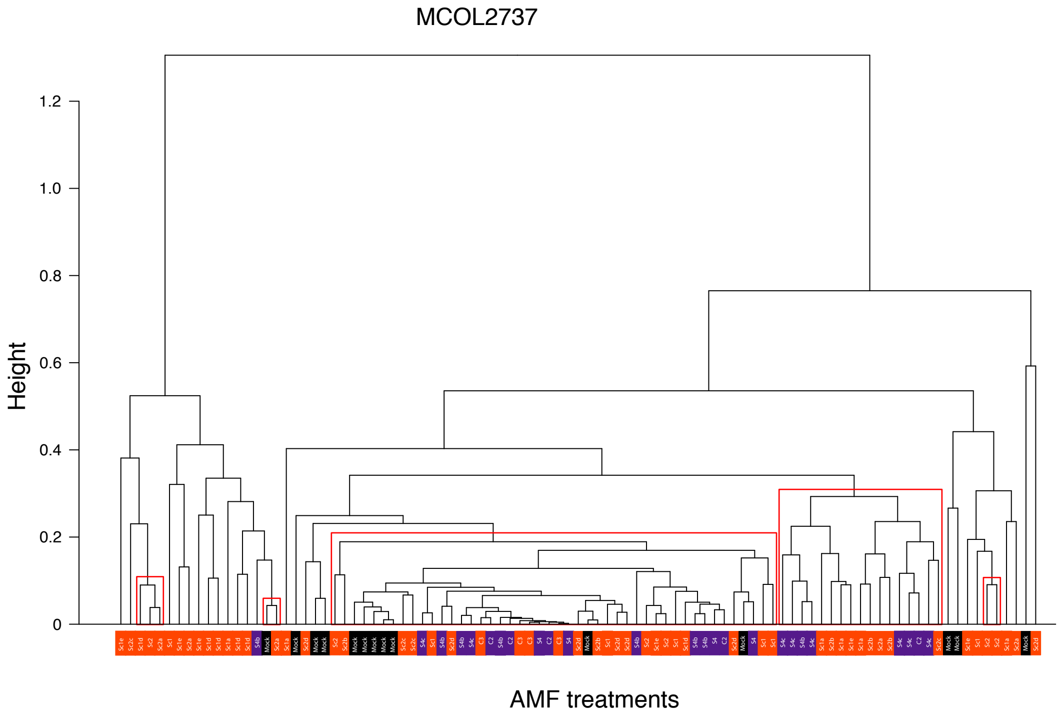


**b.**

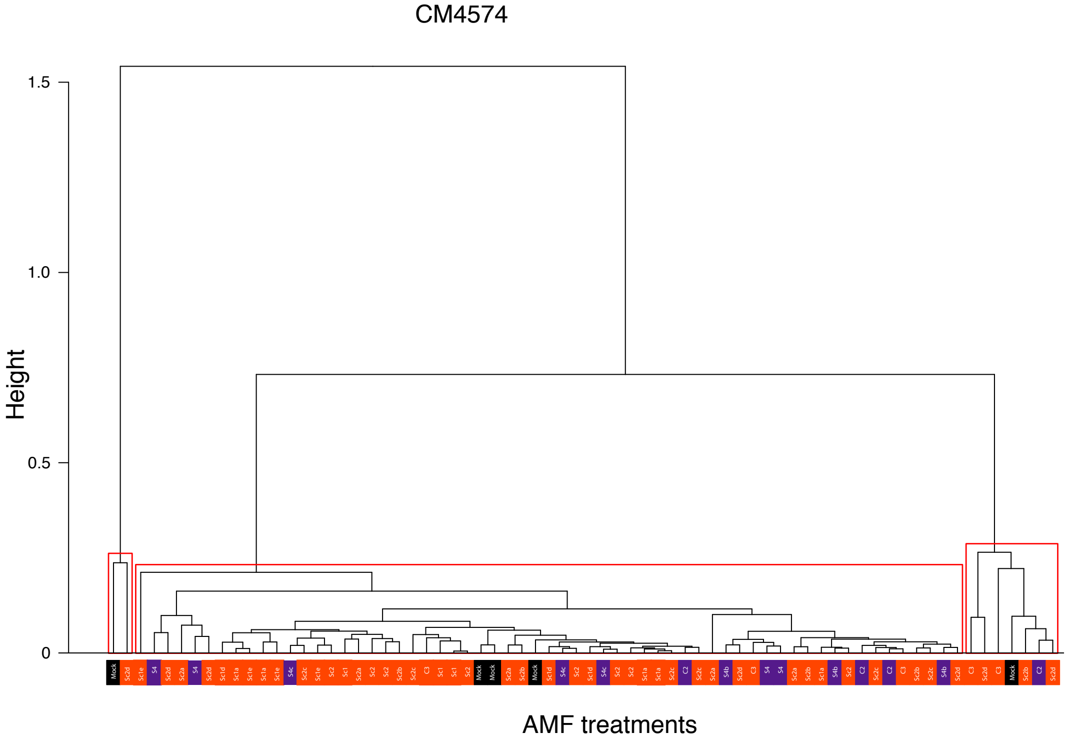

**Figure S5:** Bar-plot showing the number of reads obtained from the same plants sampled at the 3 and 12 months following inoculation. There was no significant difference (see Table S8).

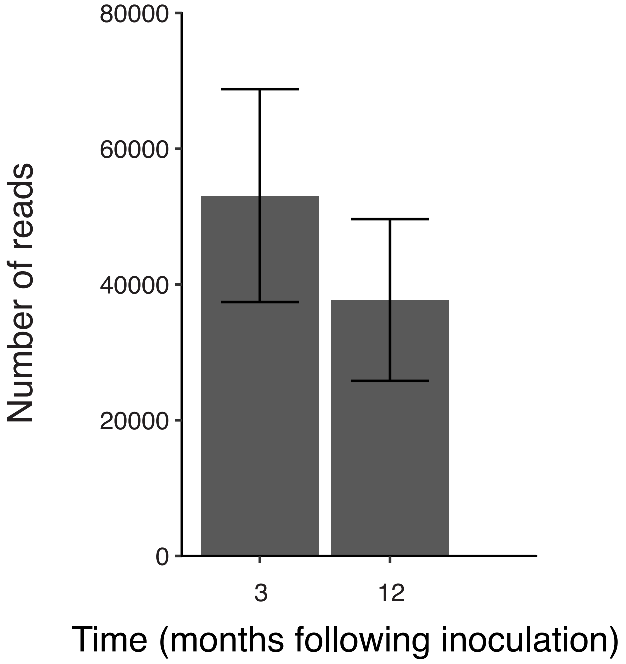


**Figure S6:** Bar-plot showing the number of OTUs obtained from the same plants sampled at the 3 and 12 months following inoculation. There was no significant difference (see table S8).

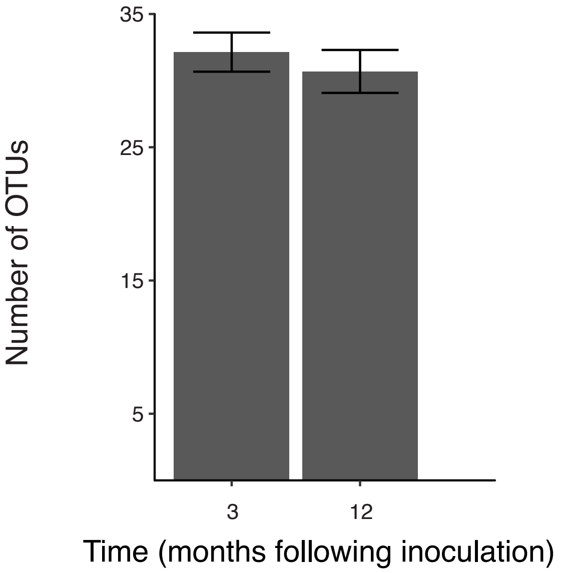
