## Supplementary note 1: Pipeline for OTU model selection for "Inoculation with highly-related mycorrhizal fungal siblings, and their interaction with plant genoptypes, strongly shapes tropical mycorrhizal fungal community structure"

### Supplementary note 1: Pipeline used to choose the model for optimal definition of OTUs using a hierarchical clustering approach

#### Methods

*OTU model selection*

The collection of alleles, weighted by their occurrence in the dataset, regardless of their taxonomic assignment, were assigned to OTUs using DBC454 v1.4.5 (Pagni et al 2013). The algorithm implemented in DBC454 progressively clusters reads depending on two parameters, which are the squared Euclidian distance cut-off value used to test for a clustering event and the minimum number of reads required to keep a cluster at each cut-off (Pagni et al 2013). The program validates a cluster if a fusion event occurs between two clusters, kept at the previous step, at a given dissimilarity cut-off. Thus, the formation of clusters is highly dependent on the initial and the final cut off distance chosen before running the algorithm. We, therefore, ran all possible DBC454 models between the smallest initial cut off distance available in the software and a final cut off distance of 225, which represents the inclusion of the collection of alleles into a single OTU, and a fixed minimum seed cluster size of 100 alleles. Allele taxonomic information obtained from BLAST results were used to apply a primary clustering model selection based on OTU taxonomic coherence. Therefore, we selected DBC454 models where no OTUs were composed of a mixture of AMF and non-AMF allele, or a mixture of valid AMF alleles from different families, as well as models where more or less than 5% of valid AMF alleles were rejected as noise. This selection procedure allowed us to determine the upper limit of the initial and final cut off distances in order to keep the resulting OTUs taxonomically coherent. In addition, selected models were compared with the VT classification using the Adjusted Rand Index (ARI) (Hubert and Arabie 1985) to quantify the differences between the approach we used and a commonly used AMF OTU selection approach involving the use of VT. Since selected models inferred different OTU richness and OTU partitioning in the dataset, we compared the robustness of each model to sampling bias by using Chao1 Index based on the number of reads in each replicate, as well as differences in number of AMF reads rejected in the noise, the OTU richness and read evenness. The OTU model at the optimum between the highest sampling effort and read partitioning was kept for subsequent analyses on OTU diversity and composition patterns.

### Results

#### *Clustering model selection*

The upper limit of the clustering model acceptance, as defined by the final cut-off distance, was influenced by taxonomic classification of the reads using BLAST references. A small proportion of reads with a best match reference in the Paraglomeraceae clade clustered with non-AMF alleles early in the clustering process. When these reads were reclassified as non-AMF, the upper Euclidian distance limit reached a value that met the criteria of acceptance of the number of AMF alleles assigned to a cluster (Figures A1 and A2) This allowed us to determine the final cuff-off distance.

**A1**

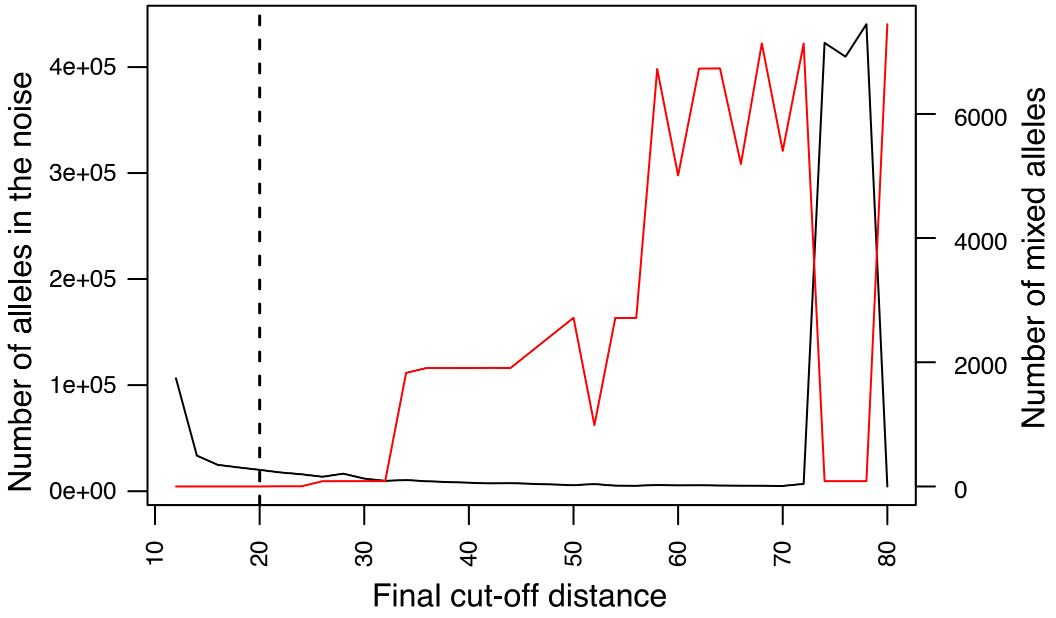

**A2**

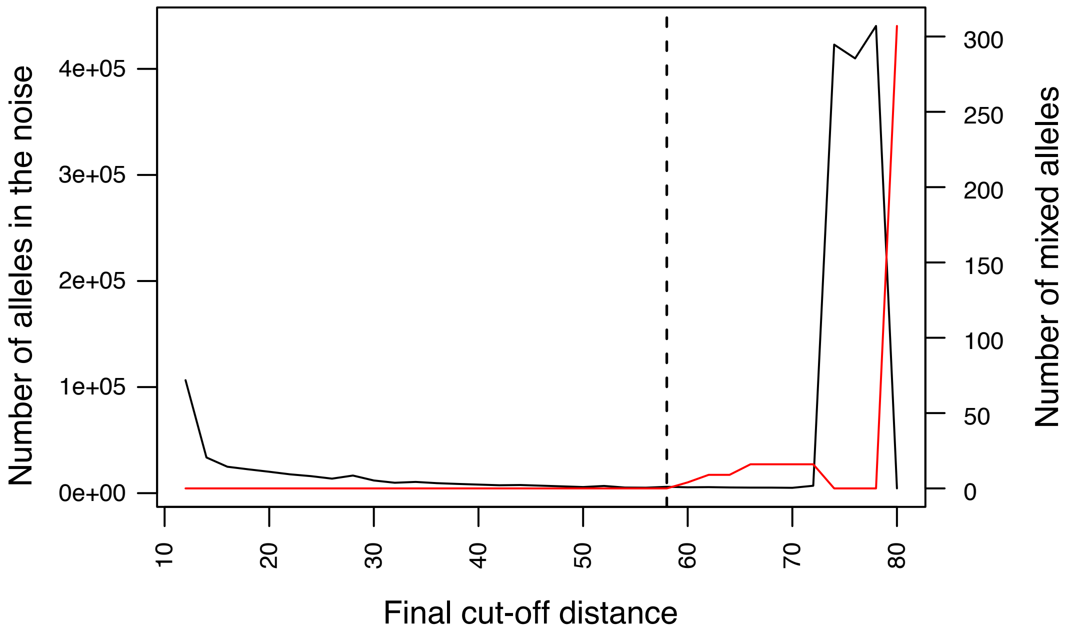


**Figure A**: Number of AMF alleles rejected in the noise (black) and number of AMF alleles mixed with non-AMF alleles in a given cluster (red) as a function of the final cut-off distance used in the DBC454 models. Results are computed based on DBC454 models using a fixed inital cut-off distance of 10, fixed cluster size of 100 and varying final cut off distance. The dashed line indicates the first final cut-off distance where the number of mixed AMF alleles was above zero. Alleles attributed to Paraglomeraceae given BLAST results are classified either as AMF **(A1)** or non-AMF **(A2).**

The lower cut-off distance selection, based on mixed clusters, showed that any model with an initial cut-off distance of between 10 and 30 to be acceptable (Figure B).

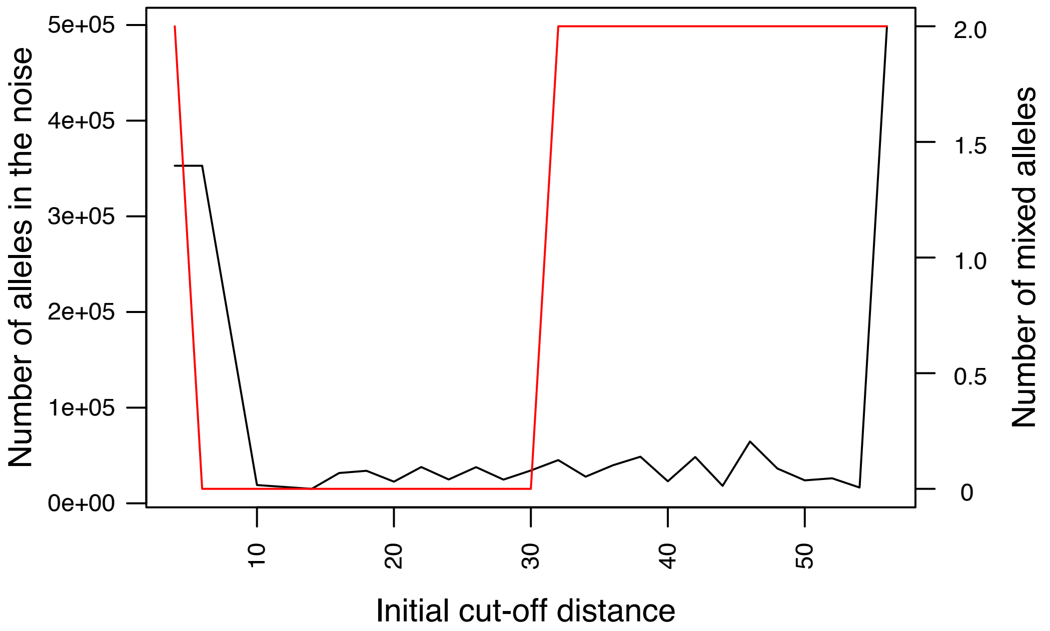


**Figure B**: Number of AMF alleles rejected in the noise (black) and number of AMF alleles mixed with non-AMF alleles in a given cluster (red) as a function of the initial cut-off distance used in the DBC454 models. Results are computed based on DBC454 models using a fixed final cut-off distance of 58, a fixed cluster size of 100 and varying inital cut-off distance.

Models with a small initial cut-off distance created more clusters with higher read partitioning and less AMF reads rejected as noise (Figure C), as well as a more similar read classification to VT defined in the MaarjAM database (Figure D).

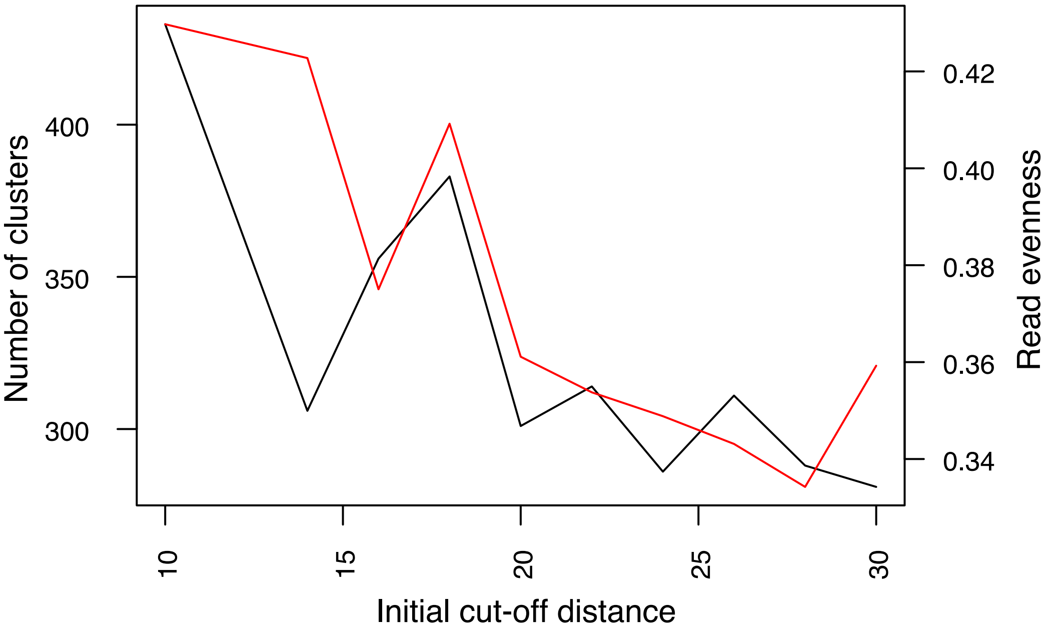


**Figure C**: Number of clusters formed (black) and read evenness index (red) as a function of the initial cut-off distance used in DBC454 models. Results are computed based on DBC454 models using a fixed final cut-off distance of 58, a fixed cluster size of 100 and varying initial cut-off distance.

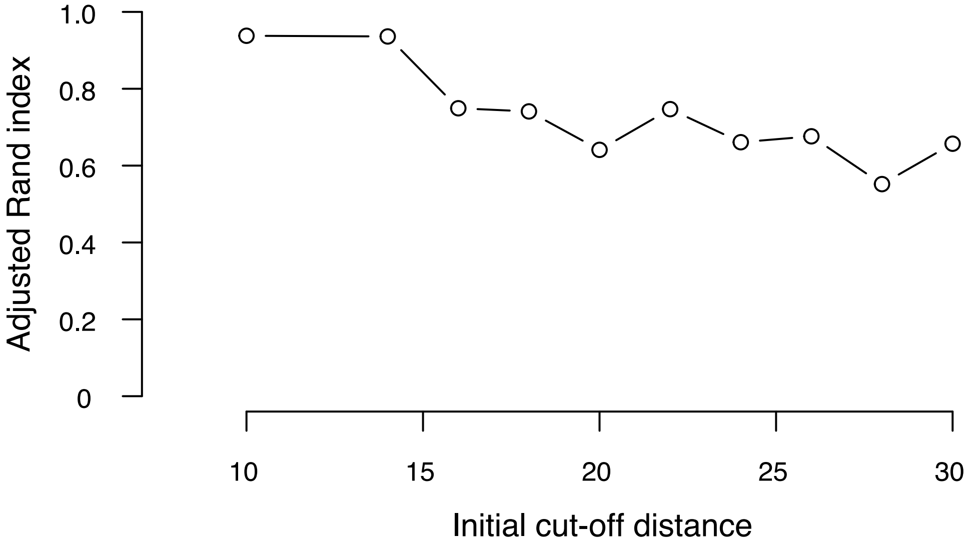


**Figure D**: Adjusted Rand index between DBC454 model and Virtual Taxa classification as a function of the initial cut-off distance used in DBC454 models. Results are computed based on DBC454 models using a fixed final cut-off distance of 58, a fixed cluster size of 100 and varying initial cut-off distance.

However, a lower cut-off distance also created more OTUs that only contained a single read per root sample (Figures E1 and E2) and, thus, the deviance between observed and estimated OTU richness increased at a low cut-off distance (Figure F).Therefore, we preferred a model that maximized sampling effort by using the clustering model with the highest acceptable initial cut-off distance (i.e., 30). Consequently, after model selection, 6092637 reads representing 535603 alleles parsed into 54 OTUs were kept for further analyses.

**E1**

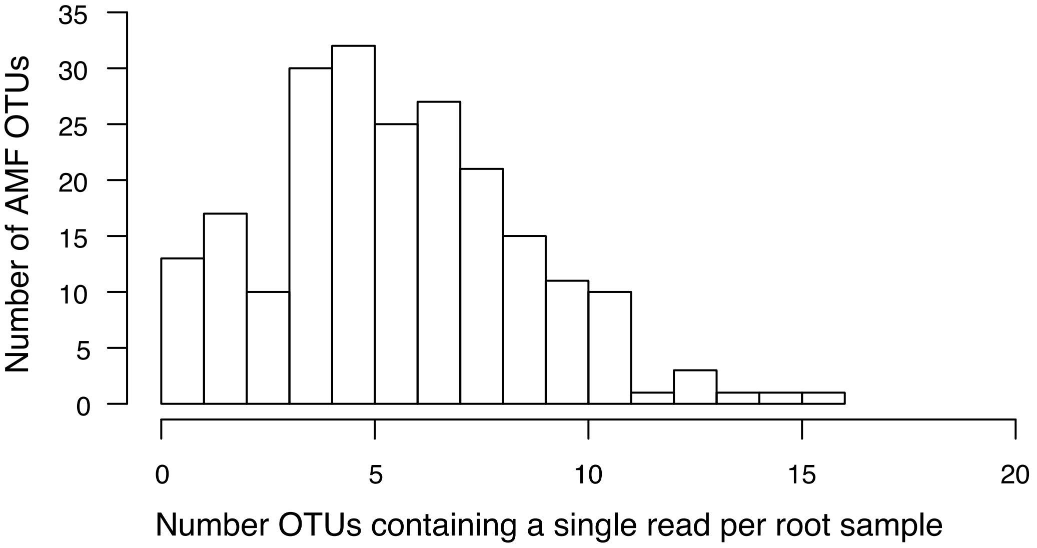


**E2**

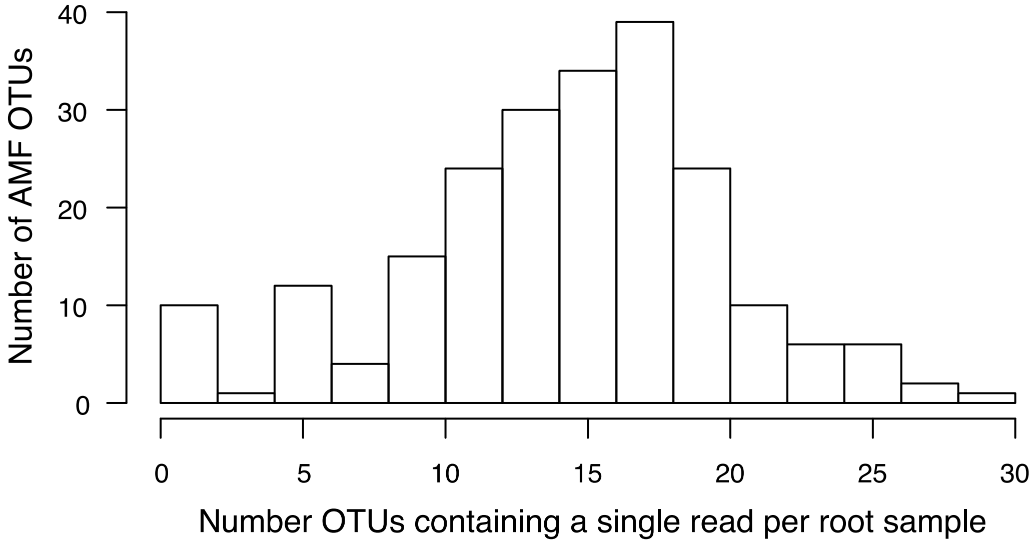


**Figure E**: Number of AMF OTUs that were represented by a single read in each root sample for **(E1)** DBC454 model with an initial cut-off distance of 10 and **(E2)** an initial cut-off distance of 30. Results are computed based on DBC454 models using a fixed final cut-off distance of 58, a fixed cluster size of 100.

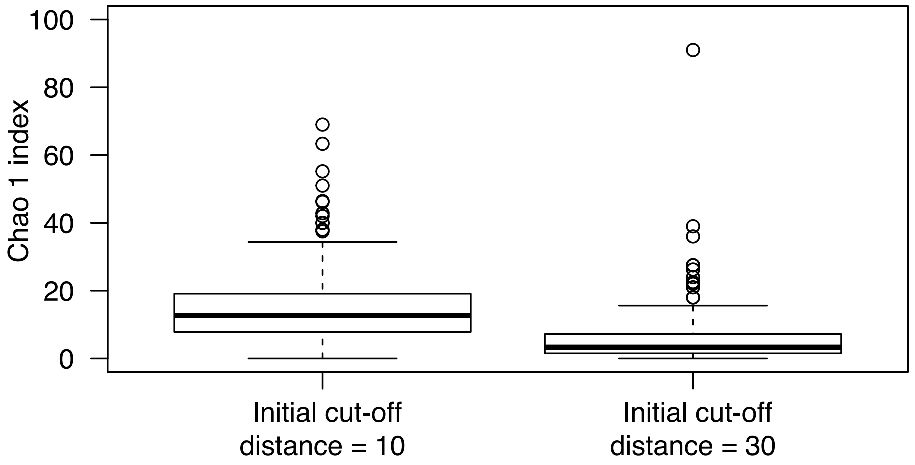


**Figure F**: Box-plots representing the difference between observed OTU richness and estimated OTU richness as given by Chao1 Index for two DBC454 models with an initial cut off distance of 10 and 30 respectively.

*Characteristics of the dataset used for analysis*

The number of AMF reads obtained from each library was highly variable, with three libraries containing less than 25% of the average read count (Figure G). However, the number of OTUs, as defined by the clustering model, was fairly equal between libraries, except for one library M04. Subsequent re-sequencing of this library (M04R) resulted in a more representative number of OTUs (Figure H).

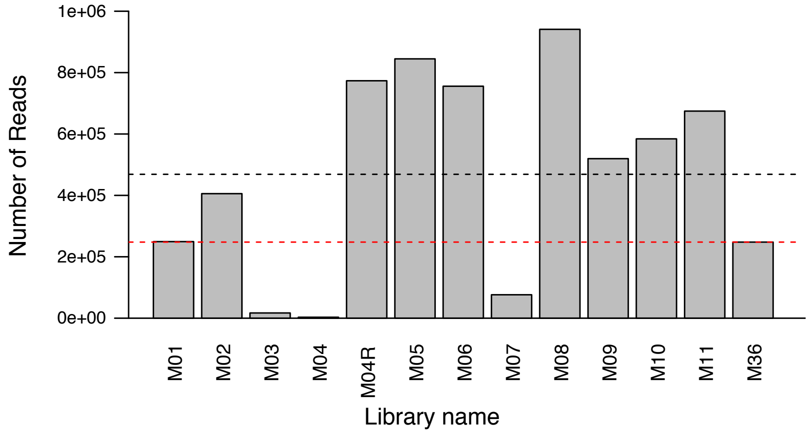


**Figure G**: Bar-plots representing the number of AMF reads obtained in each library. Black and red dashed lines represent the average and low quartile of the number of reads in the whole dataset respectively. Library M04R is a re-sequencing of the library M04

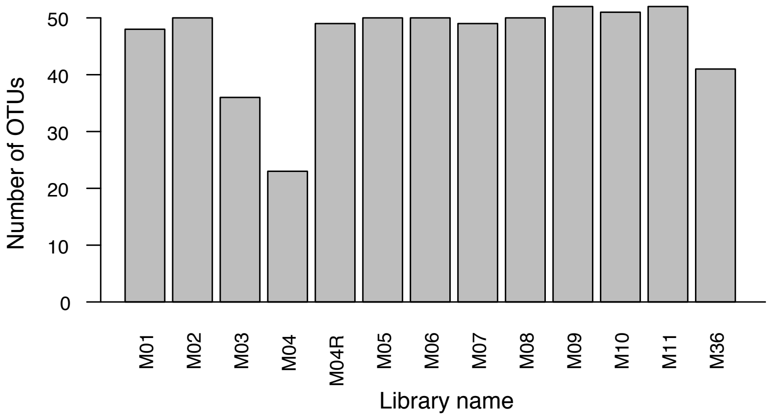


**Figure H**: Bar-plots representing the number of AMF OTUs obtained in each library. Library M04R is a resequencing of the library M04.

The number of reads per OTU was unbalanced, with one OTU encompassing more than 80% of the total number of reads in the dataset (Figure I).

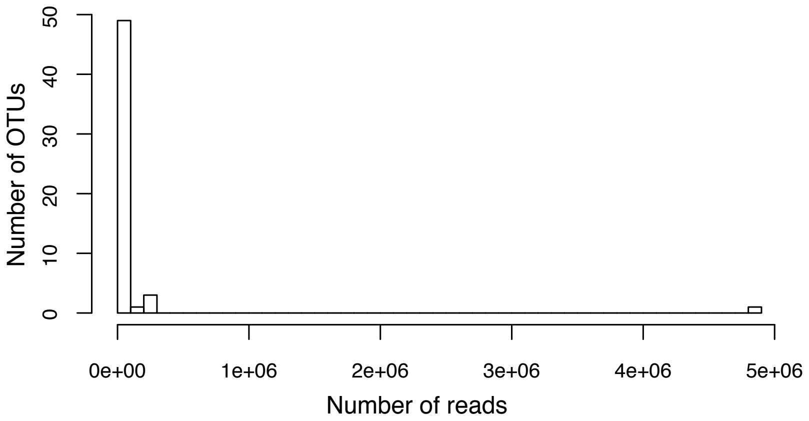


**Figure I**: Histogram showing the distribution of the number of reads per OTU.

The number of reads per root sample was also unbalanced, ranging from 19 to 400000 reads (Figure J).

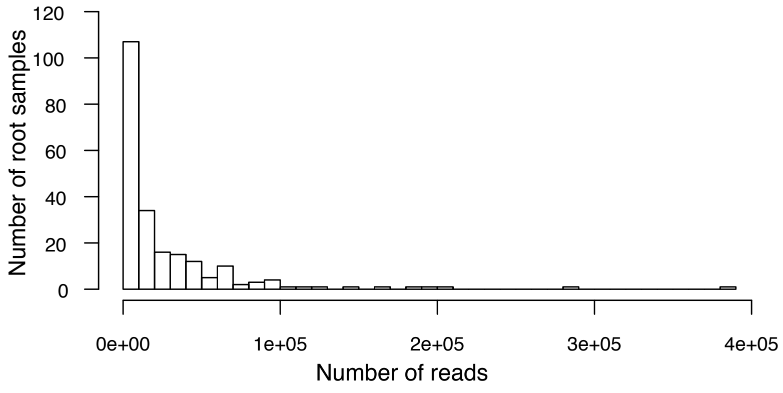


**Figure J**: Histogram showing the distribution of the number of reads per root sample.

However, the number of OTUs detected in each root sample showed a quasi-normal distribution (Figure K).

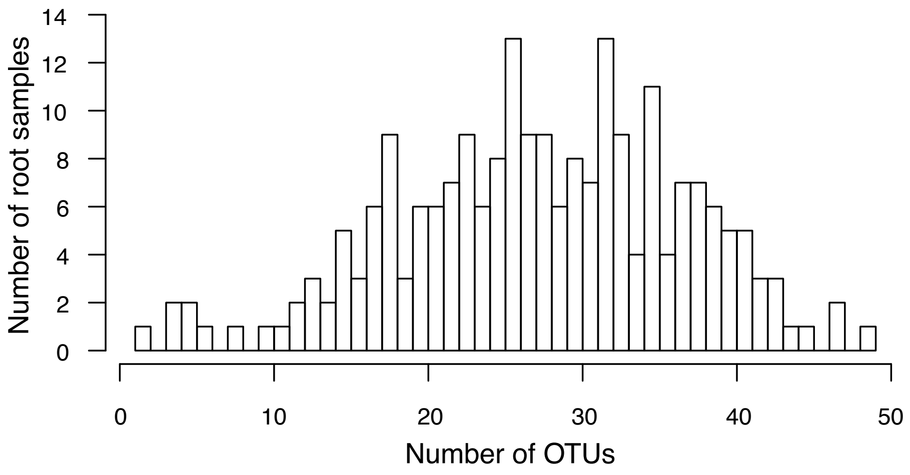


**Figure K**: Histogram showing the distribution of the number of OTUs per root sample. Distribution is very close to a normal curve (Shapiro-Wilk normality test: W = 0.9885, p-value = 0.078)

Moreover, the spatial partitioning of the OTUs in the field showed no deviance from a random spatial distribution (Figure L).

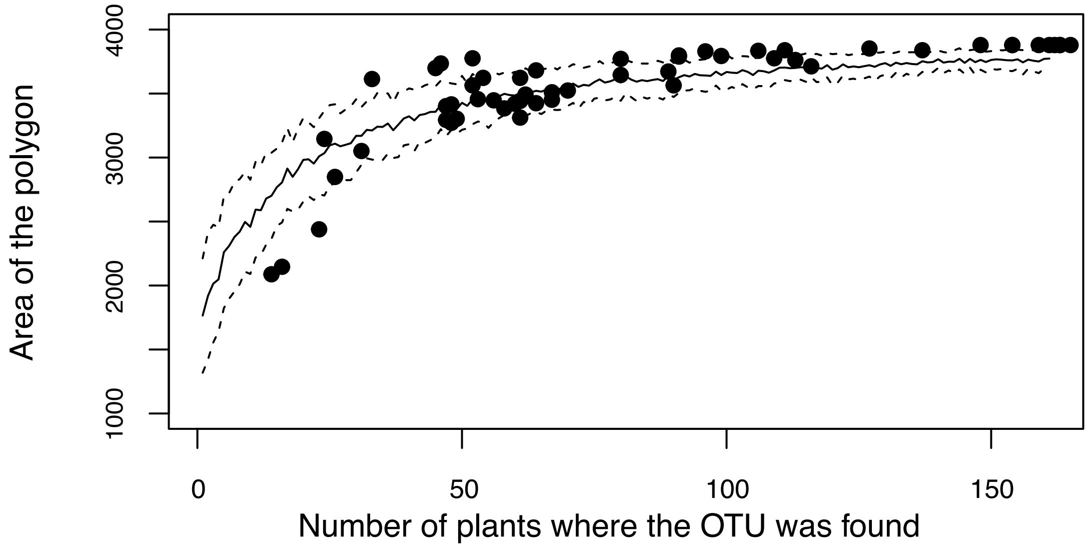


**Figure L**: Relationship between the number of cassava plants where an OTU was detected and the area of the smallest spatial polygon encompassing all the cassava plants where the OTU was detected in the field. Solid and dashed lines represent mean, 5% and 95% percentile of the null model. See text for further details about null model computation.

The number of OTUs that contained a single read per root sample was negatively related to number of reads and positively related to number of OTUs in root samples (Figure M1 and M2, respectively).

**M1**

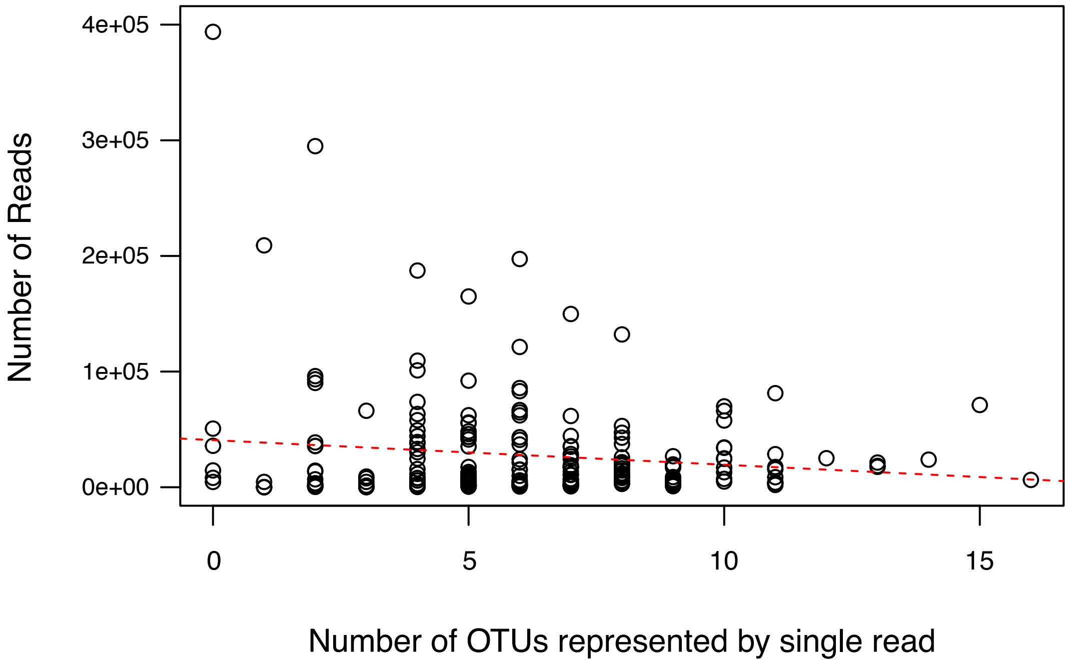


**M2**

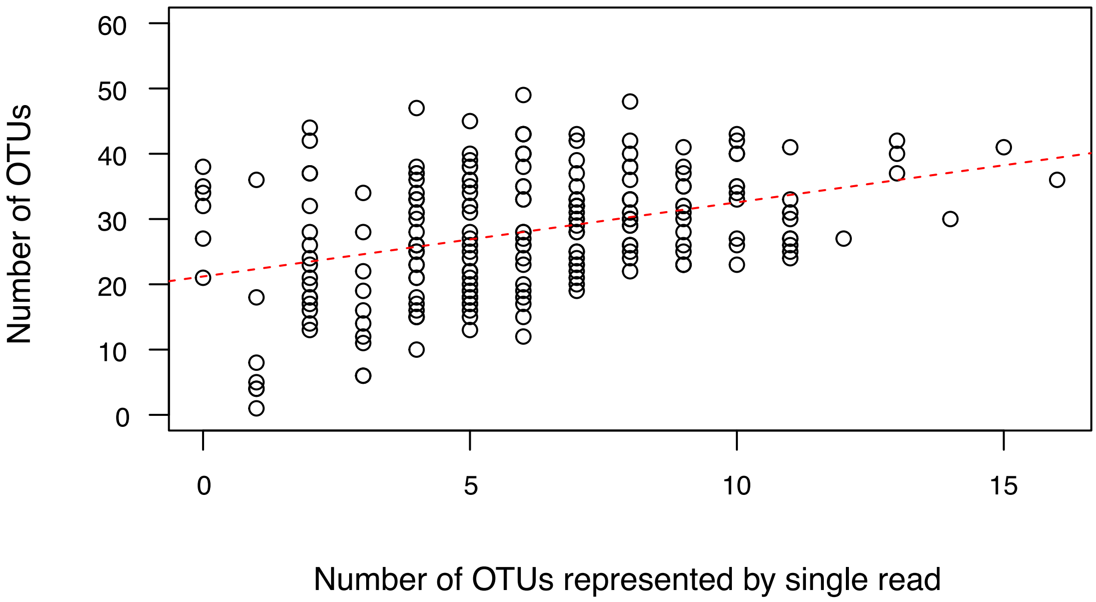


**Figure M:** Dot-plots showing the relationship between number of OTUs which contained one read and **(M1)** number of reads and **(M2)** number of OTUs. Dashed red line represents the relationship between the two variables.
